# pH-dependent anti-TfR1 NANOBODY® molecules deliver efficacious oligonucleotide payloads to muscle and CNS tissues

**DOI:** 10.64898/2026.07.29.741483

**Authors:** EA Hoyt, K Moonens, C Rapisarda, AM Salvador, TR Hammond, S Kharade, R Mondragon-Gonzalez, F Moran, S Ramkumar, J Thummapudi, S Zhou, G Haussy, C Capdevila, F Maillard, A Ismail, L Avery, P Sardi, C Nonne, S Cornelis, N Leksa

## Abstract

The blood-brain barrier (BBB) is a highly selective, semi-permeable border of endothelial cells that prevents solutes and therapeutic agents in systemic circulation from passively crossing into the central nervous system (CNS) parenchyma.

The Transferrin receptor 1 (TfR1) endocytosis pathway for iron homeostasis is one of the most well-characterized strategies for therapeutic delivery across the BBB. The work presented here showcases the discovery of novel anti-TfR1 NANOBODY® shuttles. The identified anti-TfR1 NANOBODY® molecules display cross-reactivity and pH-dependent binding to human, cynomolgus (cyno), and mouse TfR1. Structural data further explain and support the underlying mechanism of this pH-dependent binding.

These anti-TfR1 NANOBODY® molecules were successfully conjugated to both short-interfering RNA (siRNA) and antisense oligonucleotide (ASO) tool payloads. anti-TfR1 NANOBODY®-siRNA conjugates can induce up to 60% knockdown of the target mRNA transcript in skeletal muscle up to two weeks post a single IV dose in mice and up to 35-40% at four weeks post dose. Furthermore, extending the half-life of the anti-TfR1 NANOBODY®-ASO shuttles enhances heart, sciatic nerve, and brain exposure and enables up to 30-60% target knockdown in different CNS cell types.

Altogether, these results highlight important features for the development of anti-TfR1 shuttles for the purpose of downregulating target mRNA transcripts in muscle and CNS for a variety of neurologic and neuromuscular indications.

## Introduction

Strategies for tissue targeted delivery of therapeutic payloads have recently gained recognition due to their ability to significantly enhance therapeutic exposure in tissues previously considered difficult to target, such as the muscle or the central nervous system (CNS). This has repeatedly been observed for oligonucleotide-based therapeutics, for which systemic administration fails to achieve a therapeutic effect in the target tissue of interest^1^. It must be noted that oligonucleotides, such as short-interfering RNA (siRNA) and antisense oligonucleotides (ASOs), have unique chemical properties which lead to a short plasma half-life, making it difficult for them to reach the targeted tissues^2,3^. Additionally, when targeting the CNS, the oligonucleotides need to cross the blood-brain barrier (BBB), which prevents therapeutic agents from non-selectively crossing into the CNS parenchyma.

With the goal of enhancing the tissue concentration of therapeutic payloads, the Transferrin receptor 1 (TfR1) endocytosis pathway for iron homeostasis is one of the most characterized strategies for therapeutic delivery into peripheral tissues as well as the CNS. We and others have recently shown that TfR1 binding moieties can successfully be utilized for delivery of enzymes^4–7^, antibodies^8^, and oligonucleotides^9,10^ to skeletal muscle and the CNS.

There is an increasing body of literature in relation to the optimal biophysical properties and formats the TfR1 binders should have to enable payload delivery to tissues. Overall, monovalent and medium-to-low affinity to TfR1 are considered the optimal properties for delivery of payloads to the CNS^11–13^. Additionally, various reports indicate that pH-dependent binding to TfR1 (i.e. strong binding at neutral pH and weak binding at acidic pH) may be favorable for enhanced dissociation and enrichment in the CNS^14,15^. This is because physiologically, the TfR1 homodimer undergoes conformational changes when transitioning from neutral to acidic pH in the endosomal compartment. These changes involve specific histidine residues in TfR1, such as His318, His475, and His684, as well as interactions with transferrin residues like His349^16^. Structural insights from crystal structures and computational models have elucidated these dynamic processes but not yet in detail.

This work presents novel anti-TfR1 NANOBODY® shuttles that take advantage of these optimal biophysical properties when targeting TfR1 to enable delivery of therapeutic oligonucleotides to muscle and CNS for the treatment of neuromuscular indications. Here, NANOBODY® refers to the single, isolated variable domain of heavy-chain-only antibodies that circulate in Camelidae.

## Materials and Methods

### Immunization, Phage Display and primary screening

Two llamas and one alpaca were immunized with recombinant human TfR1 ectodomain protein (amino acid residues 89-760; UniProt ID P02786) following standard protocols. Peripheral blood leukocyte (PBL) samples were collected and mRNA isolated. Libraries were prepared by amplifying the coding sequence for the variable heavy chain domain of the IgG2 and IgG3 fractions and subsequent sub-cloning into phagemid vector pAX212. After superinfection of the E. coli TG1 library clones with a helper phage, the selection process was carried out using recombinant biotinylated His8-AviTag human, cynomolgus (cyno) or mouse TfR1. Complexes of TfR1 and phage were captured from solution on streptavidin coated magnetic beads. After extensive washing with PBS/0.05% Polysorbate-20, bound phages were eluted by addition of 1mg/mL trypsin. Individual clones from round 1 and round 2 selections were picked.

All individual clones were grown in 96 deep well plates (1 ml volume). Expression of monovalent NANOBODY® clones was induced by adding IPTG to a final concentration of 1 mM. Periplasmic extracts were prepared by freezing the cell pellets and dissolving them in 100 µl PBS. Cell debris was removed by centrifugation.

To determine the binding capacity of the NANOBODY® clones, crude periplasmic extracts were screened using the xMAP technology (Luminex™). Different regions of MagPlex®-C MICROSPHERES beads (Luminex®) were functionalized with recombinant human, cyno, or mouse ectodomain TfR1 protein and human ectodomain TfR2 protein. A bead mix containing 2000 beads of each region was added to each well of a 384-well flat bottom Nunc plate (ThermoFisher Scientific, 262160) and periplasmic extracts added to each well. NANOBODY® proteins were detected using anti-FLAG-PE (BioLegend, 637310) and read-out for each bead region was done using the FLEXMAP 3D (Luminex™).

### Expression and purification of human, mouse and cyno TfR1 proteins

TfR1 human, mouse and cyno proteins (UniProt accession numbers P02786 and Q62351, and GenPep accession number XP_065395696.1, respectively) were expressed using plasmids containing the sequences encoding the ectodomains (ECD) with a N-terminal His-tag for human and cyno proteins (His6::C89-F760_G142S and His6::C89-F760), while the mouse ECD sequence was His6-TEV::C89-F763. HEK293FS cells were transiently transfected using 293fectin^TM^ transfection reagent according to manufacturer (Gibco) recommendations and then grown in suspension culture for 8 days at 37°C, 115 rpm, with 0.8% CO_2_ for pH regulation, in FreeStyle 293 Expression Medium (Gibco). The culture supernatant was collected and purified using immobilized metal-affinity chromatography (IMAC), followed by size-exclusion chromatography (SEC) on a HiLoad Superdex 200 PG preparative SEC column (Cytiva) in DPBS buffer. Fractions of interest were pooled, concentrated at ca. 3 mg/mL by centrifugation on Vivaspin 20 MWCO 30 kDa (Sartorius). Glycerol at a final concentration of 10% was added prior to store TfR1 proteins at −80°C. The purity, molecular weight, and sequence of TfR1 proteins were confirmed by analytical SEC, liquid chromatography/mass spectrometry (LC/MS) and sodium dodecyl sulfate polyacrylamide gel electrophoresis. Analytical SEC was performed with a Superdex 200 Increase 5/150 column (Cytiva) that was equilibrated with Dulbecco’s phosphate buffered saline (DPBS, Gibco) at room temperature on an Agilent 1290 Infinity LC system with OpenLab software. Proteins samples were deglycosylated using Protein Deglycosylation Mix II (New England Biolabs) according to supplier’s recommendations. LC/MS was then performed using a

QExactive Plus MS spectrometer coupled with a Vanquish Core HPLC system (Thermo Scientific) equipped with a MAbPac RP analytical column (Thermo Scientific), which was thermostated at 30°C and run with LC eluant mix WF2 UpS (ROMIL Ltd) as the mobile phase. Raw data reprocessing was performed using Expressionist software (Genedata).

### Expression and purification of anti-TfR1 NANOBODY® VHHs in *E. coli*

FLAG-6xHis tagged anti-TfR1 NANOBODY® clones were selected for expression and purification. Monovalent NANOBODY® molecules were cloned into an expression vector and expressed in E. Coli TG1 cells as 3xFLAG, His6-tagged proteins. E. Coli were grown in “ZYM-5052” auto-induction medium (2 hours at 37°C followed by 29 hours at 30°C). After spinning the cell culture, periplasmic extracts were prepared by freeze-thawing the pellets and resuspending in dPBS. These extracts were used as starting material for immobilized metal affinity chromatography (IMAC) using High Affinity Ni-Charged resins (Genscript, L00223) with 0.2M Na acetate pH 4 as elution buffer followed by a desalting step with PD columns with Sephadex G25 resin (GE Healthcare, 28918008).

### Expression and purification of anti-TfR1 NANOBODY® VHHs in HEK293 cells

Monovalent GGCGGS fused anti-TfR1 NANOBODY® clones with C-tag were cloned into pcDNA3.4 expression vectors and expressed in Expi293F cells (Invitrogen/ Life Technologies, A14527). Materials included for transfection included OptiMEM (Invitrogen/ Life Technologies, 31985062) and Expi293F transfection kit reagents (ExpiFectamine and Enhancers; Invitrogen/ Life Technologies, A14524). Expi293F cells were diluted to a density of 1.8 million cells/mL the day before transfection using fresh Expi293F expression media (Invitrogen/ Life technologies, A14351), and cultured overnight at 37°C, 8% CO_2_, 110-120rpm. On the transfection day, the Expi293F cells were diluted to 2.5 million cells/mL, and ExpiFectamine^TM^ was diluted in Opti-MEM^TM^ and incubated at room temperature for 5 min. For each clone, the DNA was diluted in Opti-MEM^TM^ and added to the ExpiFectamine^TM^ /Opti-MEM^TM^ solution. The final solution of DNA/ ExpiFectamine^TM^ in Opti-MEM^TM^ was incubated at room temperature for 10-20 minutes before adding into the cells. The transfected cells were cultured at 37°C and 8% CO_2_ shaking at 110-120rpm. One day post transfection, Enhancers 1 and 2 were added into the cells, and harvested 4-5 days post-transfection by spinning down at 3000rpm for 20 minutes at room temperature. The resulting supernatant was filtered and used as a starting material for purification. The NANOBODY® clones were first purified by C-tag-affinity chromatography followed by size exclusion chromatography (SEC).

Selected anti-TfR1 NANOBODY® clones were formatted as either untagged formats or untagged glycine-glycine-cysteine (GGC) fused NANOBODY® formats, either monovalent or combined with an anti-serum albumin (SA) VHH building block for half-life extension (HLE).

### Expression and purification of anti-TfR1 NANOBODY® VHHs in *Komagataella phaffii*

NANOBODY® VHHs were produced in Komagataella phaffii at 2L or 5L scale using a general fed-batch methanol-free fermentation process^17^. The temperature of the bioreactor was controlled at 30°C, dissolved oxygen at 30% and pH at 6.0 during NANOBODY® production phase. Expression of the NANOBODY® molecules was derepressed by addition of an 80% (w/w) glycerol feed for 80-96 hours at a limiting and decreasing feeding rate (at start of derepression phase: 15 g/h/L initial volume, 4.5h after start: 8 g/h/L, 9h after start: 4 g/h/L, 62h after start until end of fermentation: 2 g/h/L).

The harvest was pH adjusted to pH 7.0 + 0.2. The harvest was clarified via microfiltration to remove cells and cell debris using tangential flow filtration with a nominal molecular weight cut off of 0.2μm, Hydrosart (Sartorius 3081860702W-SW). The NANOBODY® was then purified with Protein A chromatography. The eluate was adjusted depending on the pI of the molecule to allow for subsequent binding on ion exchange chromatography. The NANOBODY® formats were further purified to remove any truncated and self-associated forms by ion exchange chromatography with elution of the target by salt gradient. Fractions of interest were then pooled and concentrated via vivaspin spin column of appropriate nominal molecular weight cutoff to allow for further purification via SEC to further purify the NANOBODY® format from residual levels of truncated and self-associated formats and to exchange into DPBS. To reduce the chance of endotoxin an optional filtration with a Mustang E filter can be performed (Cytiva MSTG25E3) and finally filtered at 0.22μm cut off prior to storage at ≤ −20°C.

### BLI based binding quantification to TfR

The TfR binding kinetic values (Ka, Koff and KD) for each anti-TfR1 NANOBODY® were determined against human, cyno and mouse TfR1 by Bio-Layer Interferometry (BLI) using an Octet® HTX system (Sartorius). Biotinylated TfR1 from each species was immobilized on streptavidin (SA) biosensors (Sartorius, 18-5019) in HBS-P+ buffer (10mM HEPES, 150mM NaCl, 0.05% P20, pH 7.4; Cytiva, BR100671) and baseline establishment (60s), the biosensors were dipped into different concentrations (ranging from 3.91nM – 1000nM) of the anti-TfR1 NANOBODY® molecules diluted into HBS-P+ buffer for the association phase (90s). This was followed by the dissociation phase (180s) in HBS-P+ buffer.

### MSD based pH-dependent binding

The binding affinity (K_D_ value) for the human TfR1 binding of the FLAG-6xHis tagged anti-TfR1 NANOBODYs was determined at neutral and low pH values (7.4 and 6) using Meso Scale Discovery (MSD) technology. A dilution series of recombinant human TfR1 ectodomain protein was incubated with a fixed concentration of purified anti-TfR1 NANOBODY® molecules in 1x PBS + 1% BSA assay buffer (either pH 7.4 or pH 6.0) for 48h at 25°C. Next, the NANOBODY-TfR1 mixtures were applied over MSD GOLD 96-well Small Spot Streptavidin SECTOR Plate (Meso Scale Discovery, L45SA-1) that were blocked using MSD Blocker A Kit (Meso Scale Discovery, R93AA-1) and coated with 1μg/mL biotinylated human ectodomain TfR1 protein. After 10 minutes of incubation, the plates were washed and detection was performed using in house Sulfo tagged (Sulfo Kit: GOLD MSD SULFO-TAG NHS Ester-Meso Scale Discovery R91AO-1) ANTI-FLAG® M2 (Sigma-Aldrich, F3165) and MSD GOLD Read Buffer (Meso Scale Discovery, R92TG-2) with a read-out on a MESO QuickPlex SQ 120 device

### Epitope binning

Epitope binning was performed for anti-TfR1 NANOBODY® molecules with BLI using an Octet® HTX system (Sartorius). Biotinylated human TfR1 at 10nM was immobilized on streptavidin (SA) biosensors (Sartorius, 18-5019) in HBS-P+ buffer (10mM HEPES, 150mM NaCl, 0.05% P20, pH 7.4; Cytiva, BR100671).

A baseline signal (60s) was established after loading of the first anti-TfR1 NANOBODY® molecule was performed to saturate TfR binding positions (300s with 400nM molecule). Another baseline step (60s) was completed followed by a second anti-TfR1 NANOBODY® molecule capture (300s with 400nM molecule). A final dissociation step (600s) was then performed. Competition was determined by the absence of a significant difference in RU level during the second anti-TfR1 NANOBODY® molecule capture step. On the other hand, a significant increase in RU level indicated that the anti-TfR1 NANOBODY® molecules targeted different epitopes.

### Competition with transferrin (Tf)

Purified anti-TfR1 NANOBODY® molecules were tested for competition against the human TfR1 ligand transferrin (Tf). EC30 concentration of recombinant biotinylated human Tf (Sigma-Aldrich, T3915) was co-incubated with a titration series of each anti-TfR1 NANOBODY® molecule before adding them to HEK293T cells that endogenously express human TfR1. After 90 minutes incubation, the cells were washed and binding of the Tf ligand was detected with streptavidin-PE (BD-Pharmingen, 554061). Cell suspensions were analyzed with iQue Screener PLUS 3 (Intellicyt) and dose response modelling as performed using 4 parameter logistic regression in GraphPad (GraphPad Software Inc.).

Schild analysis of Tf binding in presence of high concentration of NANOBODY® molecules demonstrated no significant impact on Tf binding. A control (non-TfR1 binding) NANOBODY® and a Tf blocking NANOBODY® served as a negative and positive control, respectively. In short, human transferrin was titrated out on HEK293T cells in absence and presence of EC30, 10x EC50, and 100x EC50 concentrations of NANOBODY® molecules (pre-mixing Tf and NANOBODY® clones). After 90 minutes incubation the cells were washed, binding of the Tf ligand was detected with streptavidin-PE (BD-Pharmingen, 554061) and DAPI (BD Biosciences, 564907) was used as dead stain to gate out living cells. Cell suspensions were analyzed with iQue Screener PLUS 3 (Intellicyt) and dose response modelling as performed using 4 parameter logistic regression in GraphPad (GraphPad Software Inc.).

### Cryo-EM structure determined binding interaction between NANOBODY® and human TfR1

Localization of the human TfR1 epitopes recognized by the pH sensitive NANOBODY® molecules was determined using cryogenic electron microscopy (cryo-EM). Purified hTfR1 (positions 89 to 760) was mixed with the NANOBODY® or the antibody molecules at a molar ratio of 1:1.2 and further purified using a Superdex200 3.3/300 column pre-equilibrated with 150 mM NaCl 25 mM HEPES pH 7.4 or pH 6 when appropriate. UltraAufoil® R 0.6/1 on 300 (Quantifoil) gold mesh grids were glow discharged at 22 mA for 45 seconds and 3μL of each sample at a concentration of <1 mg/mL were added to the grids and plunge frozen in liquid ethane. The raw micrographs from the grids were collected on either a Glacios or a Krios G4 microscopes (ThermoFisher) equipped with a Falcon4 or a Falcon 4i camera at 200 and 300 keV, respectively. Images in the EER format were recorded with EPU at 240,000X and 165,000X nominal magnification at a pixel size of 0.58Å or a 0.74Å and a range of defocus from −0.8 to −2.2. Dose on camera during exposure time of 4.72 seconds was 60 and 40^e-^/Å2 respectively for each microscope. Specific details for each dataset collection are summarized in Supplementary Table 1.

More than 4,000 images were taken in total for each dataset. The data analysis was conducted using Cryosparc versions 3 to 4^18^. The final high-resolution map was obtained via non-uniform refinement^19^. The dataset for Nb2 was solved using relion 4.0^20,21^. The resolution of the final reconstructions was estimated using the value at which the FSC curve fell below 0.143^22^. The key data analysis steps for each dataset with the FSC curves and the local resolution are summarized in Supplementary Figures 1-2. The cryo-EM maps were sharpened using Phenix Autosharpen or DeepEMhancer ^23,24^ and then used to fit the atomic coordinates of the TfR1 (whether human, mouse or cyno) and the NANOBODY® of interest. Alpha fold 2 was used to build the models when no experimental structure was available.^25^ The atomic coordinates underwent several rounds of manual refinement with Coot and/or ISOLDE followed by a final real space refinement in Phenix to calculate the B factors for each residue ^26–28^. The structures were visualized using ChimeraX^29^. Electrostatic potential surfaces were calculated and visualized using the Coulombic surface coloring tool in UCSF ChimeraX, with the potential thresholds set from negative (red) to positive (blue).^29^

### Anti-TfR1 NANOBODY®-oligonucleotide conjugate generation

GGC-tagged NANOBODY® formats were conjugated to Succinimidyl-4-(N-maleimidomethyl) cyclohexane-1-carboxylate (SMCC) activated oligonucleotides and the resulting conjugates purified by ion exchange chromatography. In short, reduction of GGC-NANOBODY® dimers was performed by adding a molar excess of TCEP for 3.5 hours at room temperature. The excess TCEP was removed by desalting using 2mL Zeba desalting columns (Thermo Scientific, 89889). Subsequently, molar excess of the SMCC-oligonucleotide was added. Anion Exchange (AEX) was performed to remove excess oligonucleotide and unreacted NANOBODY® formats. Selected fractions were desalted with 2mL Zebaspin columns (Thermo Scientific, 89889). Final concentration of the NANOBODY®-oligonucleotide was determined by performing a BCA assay and quality assessed by mass spectrometry and SDS-PAGE.

MALAT1 ASO:

5’-(SMCC)(NHC6)GbsCbsAbsdTsdTs(5MdC)sdTsdAsdAsdTsdAsdGs(5MdC)sAbsGbsCb-3’

Nb: LNA residues (including LNA-5MeC and LNA T/LNA-5MeU)

dN: DNA residues

(5MdC): 5-Methyl DNA C

s: phosphorothioate backbone modification

(NHC6): Aminohexyl linker

(SMCC): SMCC NHS ester

HPRT siRNA:

Sense Strand: 5’-(SMCC)(NHC6)uscsCfuAfuGfaCfuGfuAfgAfuUfuUfaUf-3’

Antisense strand: 5’-pasUfsaAfaAfuCfuAfcAfgUfcAfuAfgGfasasu-3’

n: 2’-O-methyl residues

Nf: 2’-Fluoro residues

s: phosphorothioate backbone modification

p: Phosphate

(NHC6): Aminohexyl linker

(SMCC): SMCC NHS ester

### Radiolabeled biodistribution rodent studies

To evaluate the pharmacokinetics (PK) and biodistribution (BioD) of anti-TfR1 moieties in various tissues, anti-TfR1 proteins were radiolabeled with iodine-125 (^125^I) by indirect iodination through lysine residues using N-succinimidyl 3-^125^I-iodobenzoate (^125^ISIB) reagent. The ^125^I labelled proteins were tested to confirm the retention of binding to TfR1.

For in vivo PK and BioD studies, mice were anesthetized with isoflurane (5% isoflurane, 2L/min air) and subsequently intravenously (IV) injected via the retro-orbital (RO) venous with the appropriate radiolabeled construct using a 0.3mL syringe fitted with a 29-gauge needle. For sample collection at each time point, mice were anesthetized by intraperitoneal (IP) injection of a mixture of ketamine hydrochloride (100mg/kg) and xylazine hydrochloride (10 mg/kg) and were rapidly sacrificed by exsanguination via intracardiac puncture to collect terminal blood samples. Organs of interest were harvested, rinsed with saline and weighed using a precision balance.

The concentration of radiolabelled protein in tissues was quantified in an automatic gamma counter (Hidex AMG) calibrated for ^125^I (LLOQ: 100cpm), in a channel with windows set for 15-80 keV. The radioactivity in plasma and tissue samples was expressed as percentage of the injected dose per gram (%ID/mL) and was subsequently used to calculate the absolute molar concentration to plot against time.

### Brain parenchymal biodistribution of anti-TfR1 NANOBODY® and anti-TfR1 NANOBODY®-ASO conjugates

Anti-TfR1 NANOBODY® and anti-TfR1 NANOBODY®-tool ASO conjugates were single dosed at 200 nmol/kg IV into hTfR1-KI mice and euthanized followed by transcardiac perfusion with cold PBS at 24h post-dosing. Brain hemispheres were rapidly collected and fixed in 10%NBF for 48h at 4C, followed by washes with cold PBS. Brain hemispheres were embedded in paraffin, sectioned, and stained with fluorescently tagged antibodies to visualize anti-TfR1 NANOBODY® and anti-TfR1 NANOBODY®-ASO conjugates, NeuN as neuronal marker, Endoglin as endothelial cell marker, Hoechst as nuclear marker (please see following methods section for further details on embedding, sectioning, and staining). Multicolor immunofluorescence allowed the assessment of colocation of the VHH signal with NeuN with the aim of confirming neuronal internalization of the anti-TfR1 NANOBODY® and anti-TfR1 NANOBODY®-ASO conjugates.

### Embedding, sectioning, and staining followed by immunofluorescence (IF) imaging

Mouse brain hemispheres were first incubated overnight in 20% glycerol and 2% dimethylsulfoxide. The hemispheres were embedded for sagittal sectioning in a gelatin matrix using MultiBrain^®^ Technology (NeuroScience Associates, Knoxville, TN). The block was cured with a formaldehyde solution before being frozen in 2-methylbutane with crushed dry ice on an AO 860 microtome. The block was sectioned into 30 um sagittal sections. All sections were cut through the entire brain and collected sequentially in 12 cups of Antigen Preserve solution (50:50:1, PBS pH 7.0:ethylene glycol:polyvinyl pyrrolidone). Every 12^th^ section was stained free-floating for IF. All incubation solutions used Tris buffered saline (TBS) with Triton X100 for vehicle solutions. Primary antibody staining occurred overnight at room temperature. After rinsing, fluorescently tagged secondary antibodies were applied followed by rinsing and a Hoechst counterstain. Further rinses were completed before the sections were mounted on gelatin coated glass slides and air dried. Stained slides were dehydrated in alcohol, cleared in xylene, and cover slipped.

### In vivo knock down rodent studies

Single dose target knockdown studies were performed with anti-TfR1 NANOBODY®-siHprt conjugates, dosed at 200 nmol/kg IV into hTfR1-KI mice. At the indicated time points after the single dose, mice were euthanized with CO2 and transcardially perfused with cold PBS. Tissues were harvested and processed with quantitative PCR for Hprt and Actb as house-keeping genes. Repeated dose target knockdown studies were performed with anti-TfR1 NANOBODY®-siHprt conjugates, dosed at 200 nmol/kg IV into hTfR1-KI mice, once per week for four weeks. Mice were euthanized 1 week after the last dose with CO2 and transcardially perfused with cold PBS. Tissues were harvested and processed with quantitative PCR for Hprt and Actb as house-keeping gene.

Repeated dosing target knockdown studies with anti-TfR1 NANOBODY®-Malat1 ASO conjugates were performed in hTfR1-KI mice by IV dosing four times 400nmol/kg over two weeks, on days 0, 3, 7 and 10. 3 days after the last IV dose, mice were euthanized with CO2 and transcardially perfused with cold PBS. Tissues were harvested and processed with quantitative PCR for Malat1 and Actb as house-keeping gene.

### RNA isolation from tissues, RT and qPCR

Mouse tissues were homogenized in 2.8mm ceramic bead containing 2mL tubes, with 1mL Trizol (ThermoFisher 15596018) using a bead mill homogenizer (Omni International, 19-042E). After a 5-minute incubation of the homogenate at RT, 200uL of chloroform were added per 1mL Trizol, followed by mixing by inversion 30 times. Samples were incubated for 10minutes at RT, and centrifuged at 12,000g for 15minutes at 4°C. 350uL of the upper aqueous phase were transferred to Qiagen’s Qiacube S-block, followed by RNA extraction following manufacturer’s instructions.

cDNA was generated using Applied Biosystems^TM^ High-Capacity cDNA Reverse Transcription Kit with RNase inhibitor. qPCR was run on a QuantStudio 7 Flex, using PrimeTime® Gene Expression Master Mix (IDT) and the following primers:

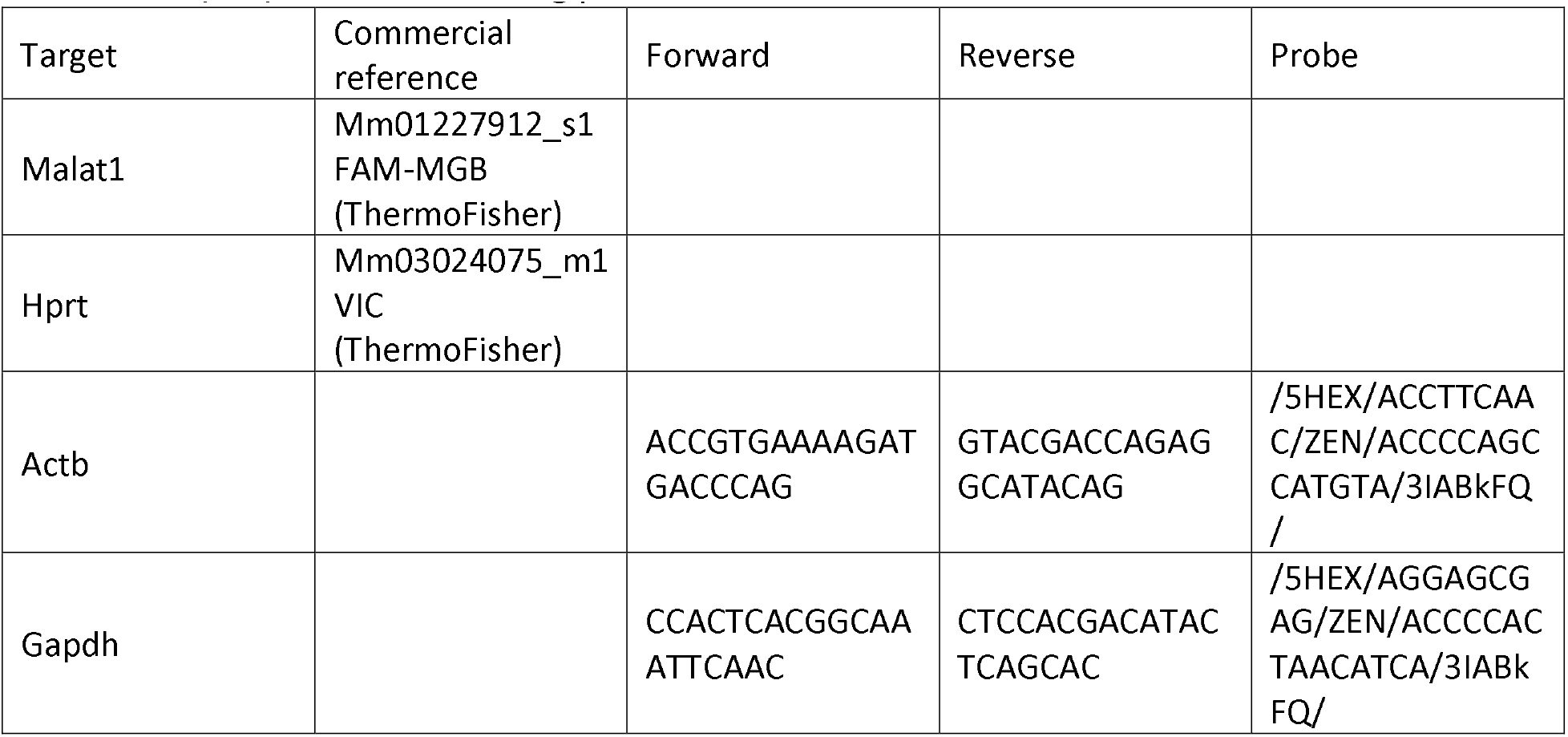

Data is represented as 2^-ΔΔCt^ as compared to saline dosed mice.

Statistical analyses were as follows: for Figure 6a-f, a 2 Way ANOVA and multiple comparisons test were performed (F value range = 0.2016-190.6; DF = 13-14); for Figure 6g, an unpaired T test versus saline was performed (t-value = 9.216, DF = 8); and for Figure 9b-f a one-way ANOVA followed by Dunnett’s multiple comparisions test versus saline was performed (F value range = 5.846-40.41; DF = 23).

### Single nucleus RNA Sequencing

Frozen mouse brain tissue or gastrocnemius muscles (15-20 mg) were processed for nuclei isolation using the Chromium Nuclei Isolation Kit with RNAase inhibitor (1000494, 10X Genomics), following manufacturer’s instructions. Nuclei suspensions were subsequently processed for single-nucleus RNA sequencing using the Chromium Next GEM Single Cell 3’ Reagents Kit v3.1 (Dual Index) (PN-1000268, 10X Genomics). Briefly, nuclei were loaded onto the Chromium Next GEM Chip G for GEM generation and barcoding. Following GEM-RT cleanup and cDNA amplification (12 cycles), 3’ Gene expression dual index libraries were constructed using 14 cycles of sample index PCR. Library quantification and QC were performed using Agilent D5000 Screentape and Reagents (5067-5589, 5067-5588, Agilent) on an Agilent

TapeStation system. Libraries were normalized to a concentration of 1.5 nM, denatured, and processed for sequencing following ‘Novaseq 6000 System denature and dilute instructions’ (Illumina), using an S2 Cluster Cartridge. Libraries were sequenced on a NovaSeq 6000 S2 flow cell using paired-end, dual indexing to a minimum of 20,000 read pairs per nucleus.

### Single nucleus RNA sequencing object setup, QC, and clustering

Samples were analyzed using Seurat 4.0.4 in R version 4.1.0. Samples were filtered to limit the inclusion of low-quality nuclei. Cells with fewer than 500 genes and 2000 UMIs were removed from analysis. To reduce ambient RNA count contamination, the SoupX package was used^30^. The object with the SoupX corrected counts was normalized and variable features were selected using the ‘vst’ method with 2,000 features. The dataset was scaled and RunPCA was performed for 60 principal components (PCs), then the FindNeighbors function was run using 40 PCs to build the k-nearest neighbor (KNN) graph. Cluster resolution was set to ‘0.1’, and the uniform manifold approximation and projection (UMAP) was generated using the RunUMAP function and 40 PCs.

Cluster identities were determined by calculating enriched markers using the FindAllMarkers function in Seurat. Cell types were assigned by identifying genes unique to each cluster and by cross-referencing known markers of each cell type from existing published datasets and with help from the scMayoMap package^31^. UMAP and gene expression plots were generated using built-in Seurat/ggplot2 (3.5.2) plotting functions or scCustomize version 2.1.2 unless otherwise described.

### Pseudobulk analysis

Pseudobulk counts matrices were generated by extracting the raw counts for a chosen cell type from the parent object and summing RNA counts for all the cells for each animal. Differential expression analysis was performed on the pseudobulk counts matrices using DESeq2 version 1.42.1 in R version 4.3.2. Normalized Malat1 gene expression counts were extracted and used for downstream plotting.

## Results

### Anti-TfR1 NANOBODY® molecules display triple species cross-reactivity

To generate anti-TfR1 NANOBODY® molecules, immunization of Camelidae family animals was performed with recombinant human TfR1 ectodomain protein. Anti-TfR1 species cross-reactive binders were enriched by performing phage display in subsequent rounds on recombinant human, cyno, or mouse TfR1. Primary screening confirmed binding to these TfR1 species and a selected panel of NANOBODY® clones (Nb1-Nb3) was produced and further characterized.

The kinetic values for each anti-TfR1 NANOBODY® molecule against the different species TfR1 were determined. Table 1 shows the kinetic values for anti-TfR1 NANOBODY® molecules binding against human, cyno and mouse TfR1 at pH 7.4. KD values indicate that all anti-TfR1 NANOBODY® molecules display a single digit nM or sub-nM affinity to hTfR1, while the affinities from cTfR1 ranged from 16 nM to 24 nM and 7.4 nM to 89nM for mTfR1. Interestingly, the affinity differences observed across species appeared to be driven by different koff values rather than ka values.

**Table 1.**
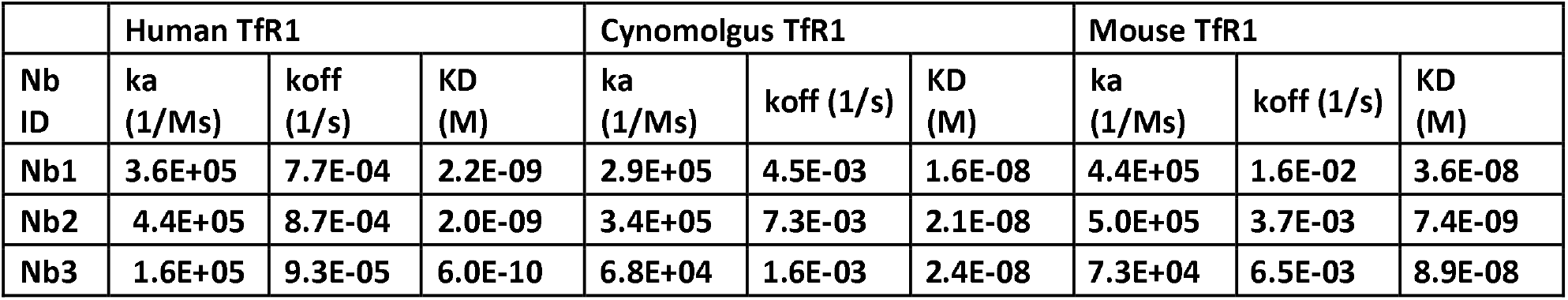
Affinity of anti-TfR1 NANOBODY® molecules to human, cynomolgus and mouse TfR1 at pH 7.4.

### Anti-TfR1 NANOBODY® clones target the same non-transferrin competing TfR1 epitope

To determine whether the anti-TfR1 NANOBODY® molecules bind the same or distinct hTfR1 epitopes, BLI binning experiments were performed. After saturation of hTfR1 with Nb2, the hTfR1-binding ability of all other anti-TfR1 NANOBODY® molecules was determined. Increase in BLI signal upon adding the second anti-TfR1 NANOBODY® indicates that the anti-TfR1 NANOBODY® molecules bind different hTfR1 epitopes; on the contrary, lack of signal enhancement indicates competition of the anti-TfR1 NANOBODY® molecules for the same TfR1 binding epitope. All anti-TfR1 NANOBODY® molecules exhibited competition for TfR1 binding, suggesting they share the same epitope bin on TfR1. All anti-TfR1 NANOBODY® molecules also showed a non-competing signal when compared to a transferrin-blocking Nb (Figure 1a).

**Figure 1.**
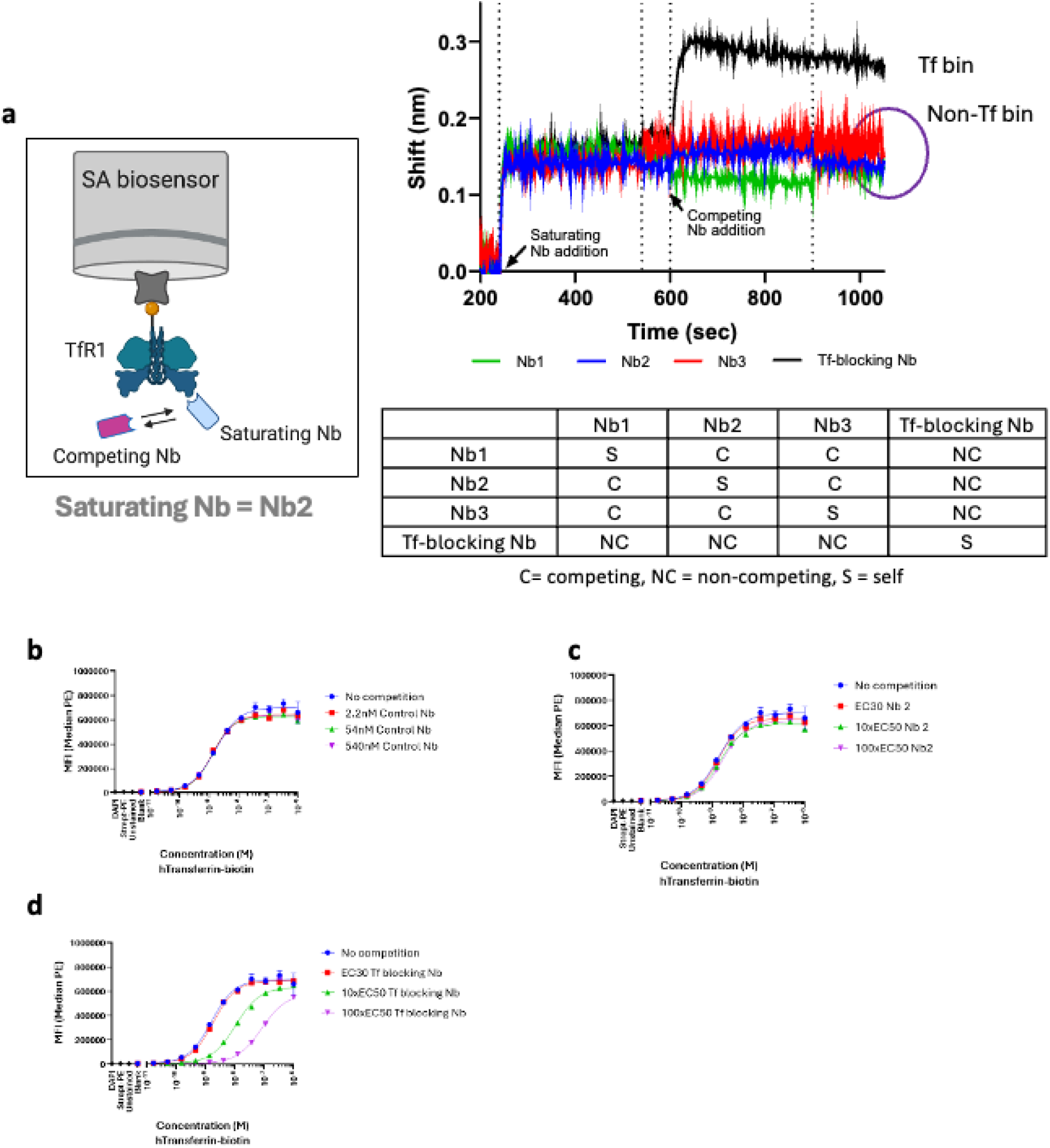
Human transferrin (Tf) competition and epitope binning exhibit the ability of anti- TfR1 NANOBODY® clones to bind hTfR1 in the presence of human Tf. (a) Epitope binning method setup and results when binning Nb1-Nb3 and a Tf-blocking Nb control against Nb1. (b-d) Schild analysis of anti-TfR1 NANOBODY® clones. Human Tf is titrated out on HEK293T cells in absence and presence of EC30, 10xEC50, and 100xEC50 concentrations of NANOBODY® protein. Results are shown for a Tf blocking Nb, Nb2, and a control Nb (with no TfR1 binding).

To further refine our understanding of the binding, flow cytometry competition assays were used to demonstrate that the anti-TfR1 NANOBODY® molecules do not directly compete with transferrin (Tf). When performing a Schild analysis, no significant shifts in EC50 value of the human Tf binding to TfR1 were observed in presence of either 10x or 100x EC50 of Nb 2 (1.3-fold difference when working in presence of 100x EC50 Nb 2). On the contrary, significant shifts in EC50 value were noticed for the Tf-blocking control Nb (respectively about 6-fold and 50-fold shifts to weaker Tf-binding in presence of 10x or 100x EC50 of the Tf-blocking Nb control). A non-TfR1 binding control Nb was also included in the assessment and, as expected, did not impact Tf binding (Figures 1b-1d). As Nb1-Nb3 share the same TfR1 epitope, Schild analysis would be expected to be comparable across all three molecules.

### Identification of the binding site of the anti-TfR1 NANOBODY® molecules by cryo-EM

We determined the cryo-EM structures of three NANOBODY® molecules in complex with hTfR1, achieving resolutions below 3 Å for all the complexes. Structural analysis revealed that all three NANOBODY® molecules recognize overlapping epitopes primarily located at the apical helical domain of hTfR1, with additional contacts extending to the protease domain of an adjacent subunit (Figure 2a-f). The NANOBODY®-hTfR1 interfaces are stabilized by a network of electrostatic interactions, including salt bridges, hydrogen bonds, and van der Waals contacts (Figure 2 g-i).

**Figure 2.**
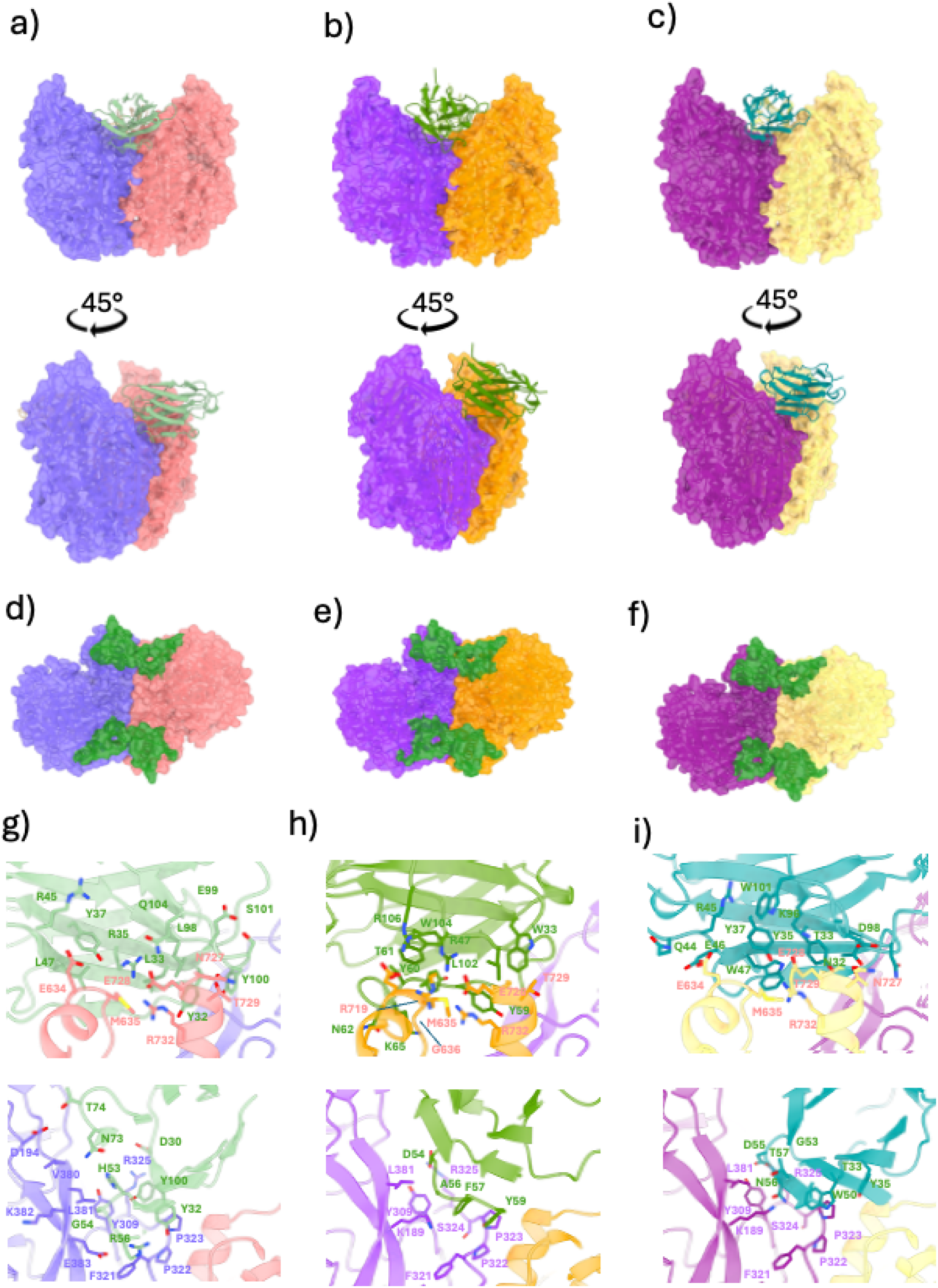
Epitope mapping reveals overlapping binding sites and distinct interaction patterns for anti-TfR1 NANOBODY® molecules. Structural characterization of NANOBODY® binding epitopes on the transferrin receptor. (a-c) Side view of the TfR1 structure showing the binding epitopes (highlighted) for NANOBODY® molecules Nb1, Nb2, and Nb3, respectively. (d-f) Top view of the isolated epitope regions (shown in green) for Nb1, Nb2, and Nb3, respectively, illustrating the spatial extent and surface area of each NANOBODY® binding site. (g-i) Detailed view of the epitope-paratope interface residues for Nb1, Nb2, and Nb3, respectively. Residues are color-coded by chain: violet hues represent TfR1 A chain residues, orange-yellow hues represent TfR1 B chain residues, and green hues represent NANOBODY® residues involved in the interaction.

The epitope of the three NANOBODY® molecules is well conserved across the three species (Supplementary Figure 3a-b), which explains the cross-reactivity of molecules with TfR1 from human, mouse and cyno (Table 1). One mutation of an Arginine to a Glutamine, in both cyno and mouse (Supplementary Figure 3c), might explain the lower affinity of the NANOBODY® molecules for the latter species compared to human (Table 1). A second mutation from KE to RD in mouse only should not affect the binding, as it is only the backbone of these amino acids that interacts with the NANOBODY® molecules (Supplementary Figure 3d). Furthermore, the CDR1 of Nb3 extends toward glycosylation site Asn727, where it engages the branched N-glycan moiety (Supplementary Figure 4a). The lower affinity of the mouse TfR1 for Nb3 (Table 1) may be further attributed to differences in glycosylation at position Asn727, which is either absent or exhibits reduced glycan occupancy in the mouse ortholog (Supplementary Figure 4b). To note, Nb2 exhibits a kinked CDR3 structure due to its germline, which causes its extended CDR to fold back upon itself, positioning key residues to interact with the glycosylation site at Asn727 but not with the glycan itself (Supplementary Figure 4c).^32^

### Anti-TfR1 NANOBODY® molecules show pH-dependent binding towards TfR1

As the co-complex cryo-EM structures highlighted, our anti-TfR1 NANOBODY® clones bind to the TfR1 dimerization interface, contacting both monomers of the TfR1 dimer. Therefore, we speculated if a conformational change in the TfR1 dimer associated with endosomal acidification could impact their TfR1 binding capacity. An in-solution affinity determination at pH 7.4, reflecting the extracellular binding to TfR1, and pH 6.0, reflecting the binding in the endosomal compartment, was performed (Figure 3). Remarkably, all tested clones showed a pH-dependent mode of binding with reduced binding at acidic pH, with the ratio of binding at pH 6.0 versus pH 7.4 varying up to 250-fold. An anti-TfR1 clone targeting the apical domain, which is distant from the TfR1 dimerization interface, was included in the analysis and, as expected, did not demonstrate the pH-dependent mode of binding (Figure 3). From a functional perspective, there is a potential benefit of having improved binding at the extracellular over the endosomal compartment as it promotes the NANOBODY® dissociation and release rate after TfR1 mediated uptake and transcytosis to the CNS.

**Figure 3.**
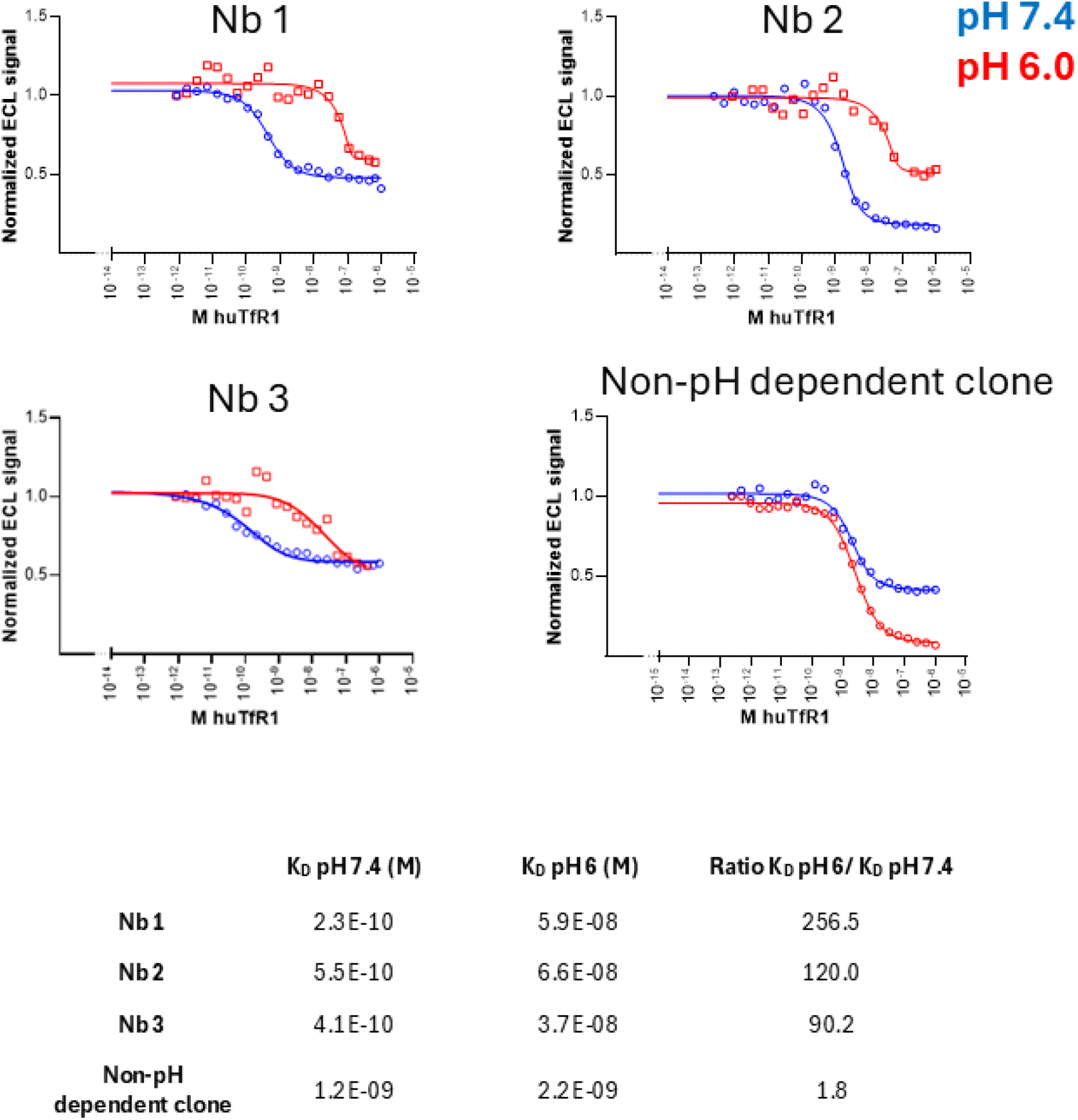
pH dependent binding of NANOBODY® molecules targeting the dimerization interface. MSD affinity determination of anti-TfR1 NANOBODY® molecules towards recombinant hTfR1 at pH 7.4 or 6.0. All identified species cross-reactive clones that target a similar TfR1 epitope exhibit a decreased affinity at pH 6. As a control, a TfR1 binding clone that targets the apical domain was included in the comparison and did not show pH-dependent binding. The affinities are summarized in the table in the lower panel of the figure.

### pH-dependent conformational changes of TfR1 drive dissociation of anti-TfR1 NANOBODY® molecules at low pH

To investigate the structural basis for the pH-dependent binding behavior of the anti-TfR1 NANOBODY® clones, we performed comparative structural analysis of the transferrin receptor at physiological pH (7.4) and acidic endosomal pH (6.0). This analysis revealed two distinct pH-dependent conformational changes, both mediated by histidine residues (Figure 4), one local and one global.

**Figure 4.**
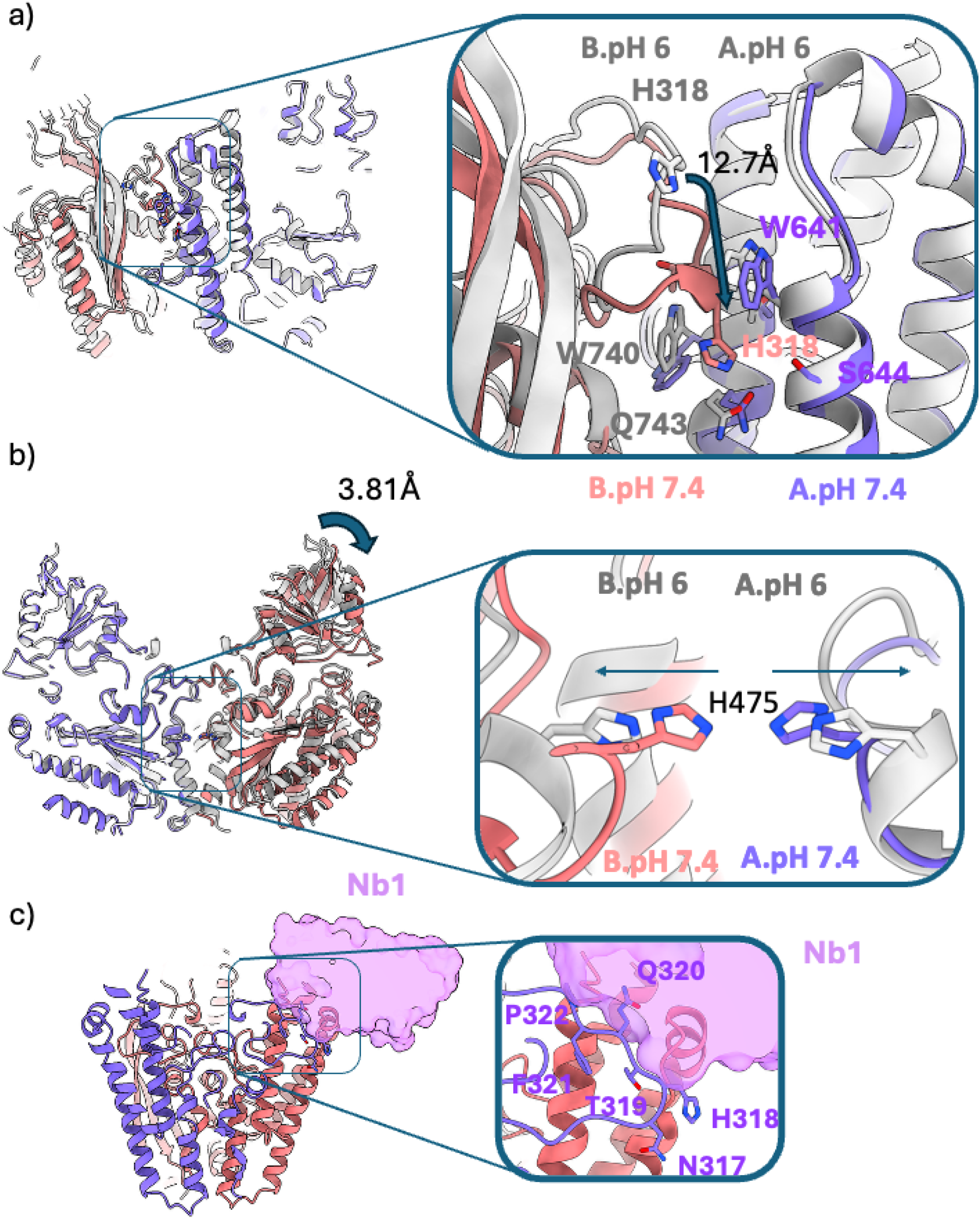
Transferrin receptor exhibits two distinct pH-dependent conformational changes involving key histidine residues. Structural analysis of TfR1 conformational transitions between endosomal (pH 6.0) and physiological (pH 7.4) pH conditions reveals two independent structural rearrangements. (a) Clipped view showing the first conformational change involving residues H318, W64, W740, and S644. At pH 6.0, protonation of H318 residues in both chains creates electrostatic repulsion, resulting in a separation distance of 12.73 Å between the two H318 residues. The disruption of favourable π-π stacking and cation-π interactions between H318 and the surrounding tryptophan residues (W64, W740), along with altered hydrogen bonding with S644, drives this conformational rearrangement. (b) Side view illustrating the second conformational change centered on His475 residues, showing the positions in both A and B chains at pH 6.0 (A.pH 6, B.pH 6) and pH 7.4 (A.pH 7.4, B.pH 7.4). At pH 6.0, protonation of the His475 residues creates electrostatic repulsion that causes a 3.81 Å displacement of the entire TfR1 structure. These two independent pH-sensitive conformational switches demonstrate the complex structural dynamics of TfR1 that facilitate its function during receptor-mediated endocytosis and iron release in acidic endosomal compartments. C) The transferrin receptor (TfR1) at pH 6 shown in ribbon representation with loop 317-321 displayed in stick format. The enlarged inset shows critical amino acid residues (N317, H318, T319, Q320, F321, P322) within a prominent surface loop of the TfR1 that creates steric clashes with the approaching NANOBODY® molecule Nb1 (shown in pink surface representation). This loop acts as a physical barrier, preventing optimal positioning and binding of the NANOBODY® molecule to its intended epitope on the receptor surface.

The local conformational change involves His318 and its interaction network with residues Trp64, W740, and S644 (Figure 4a). At physiological pH (7.4), H318 exists in its neutral state, and the loop adopts a compact conformation. Upon acidification to pH 6.0, protonation of H318 in both chains results in a separation distance of 12.73 Å between the two H318 residues. This dramatic rearrangement displaces the loop region, positioning it over the NANOBODY® binding epitope identified in Figure 2.

The second conformational change is centered on His475 residues in both the A and B chains (Figure 4b). Comparison of the structures at pH 7.4 and pH 6.0 shows that protonation of His475 at acidic pH causes a 3.81 Å displacement of the apical domain of the TfR1 structure, representing a global conformational shift.

All three characterized NANOBODY® clones (Nb1, Nb2, and Nb3) bind to TfR1 at pH 7.4 but lose binding affinity at pH 6.0 (Figure 3). Structural mapping revealed that the H318-containing loop, which undergoes the 12.73 Å displacement upon acidification, directly overlaps with the Nb binding epitope at pH 6.0 (Supplementary Movie 1). At physiological pH (7.4), the loop is retracted, leaving the epitope residues from both A and B chains (identified in Figure 2g-i) fully accessible. At pH 6.0, the loop displacement sterically occludes these same epitope residues, preventing NANOBODY® access (Figure 4c, Supplementary Movie 2). Therefore, the structural rationale behind this pH-dependent binding mode stems from the specific binding epitope on TfR1 rather than the insertion of His residues in the paratope design.

### Anti-TfR1 NANOBODY® molecules exhibit altered binding kinetics when conjugated to an ASO but not when conjugated to an siRNA

Anti-TfR1 NANOBODY® molecules were successfully conjugated to both ASO and siRNA tool payloads (Malat1 ASO and siHprt respectively) through modification of an inserted C-terminal Cys residue. Further characterization and in vitro measurement of binding kinetics at pH 7.4 between anti-TfR-NANOBODY® conjugates and hTfR1 provided evidence that the presence of the Malat1 ASO alters the interaction between the conjugate and hTfR1 (Supplementary Table 2). The BLI kinetic measurements showed a ∼10-fold or greater change to KD for Malat1 ASO conjugates while the siHprt conjugates showed no significant changes (i.e. the KD value was within 2-fold of the unconjugated KD value). Changes in binding affinity due to ASO conjugation have been reported by others as well.^33^ Due to the differences observed in the binding kinetics between the different conjugate types and the unconjugated anti-TfR1 NANOBODY® molecules, further explanation was sought through structural data.

### Structural evidence for ASO binding to the TfR1-NANOBODY® (Nb1) complex

Cryo-EM analysis of the Nb1-ASO-hTfR1 complex and the Nb1-HLE-ASO-hTfR1 revealed additional electron density that could not be attributed to either the TfR1 receptor or the NANOBODY® itself (Figure 5a and 5b, respectively). This extra density was attributed to the ASO molecule binding to the helical domain of the receptor that is usually involved in Tf binding. Meanwhile, the Nb1-siRNA-hTfR1 complex did not reveal an additional electron density, indicating that only single-stranded and not double-stranded oligonucleotides can bind to the helical domain of TfR (Figure 5c).

**Figure 5.**
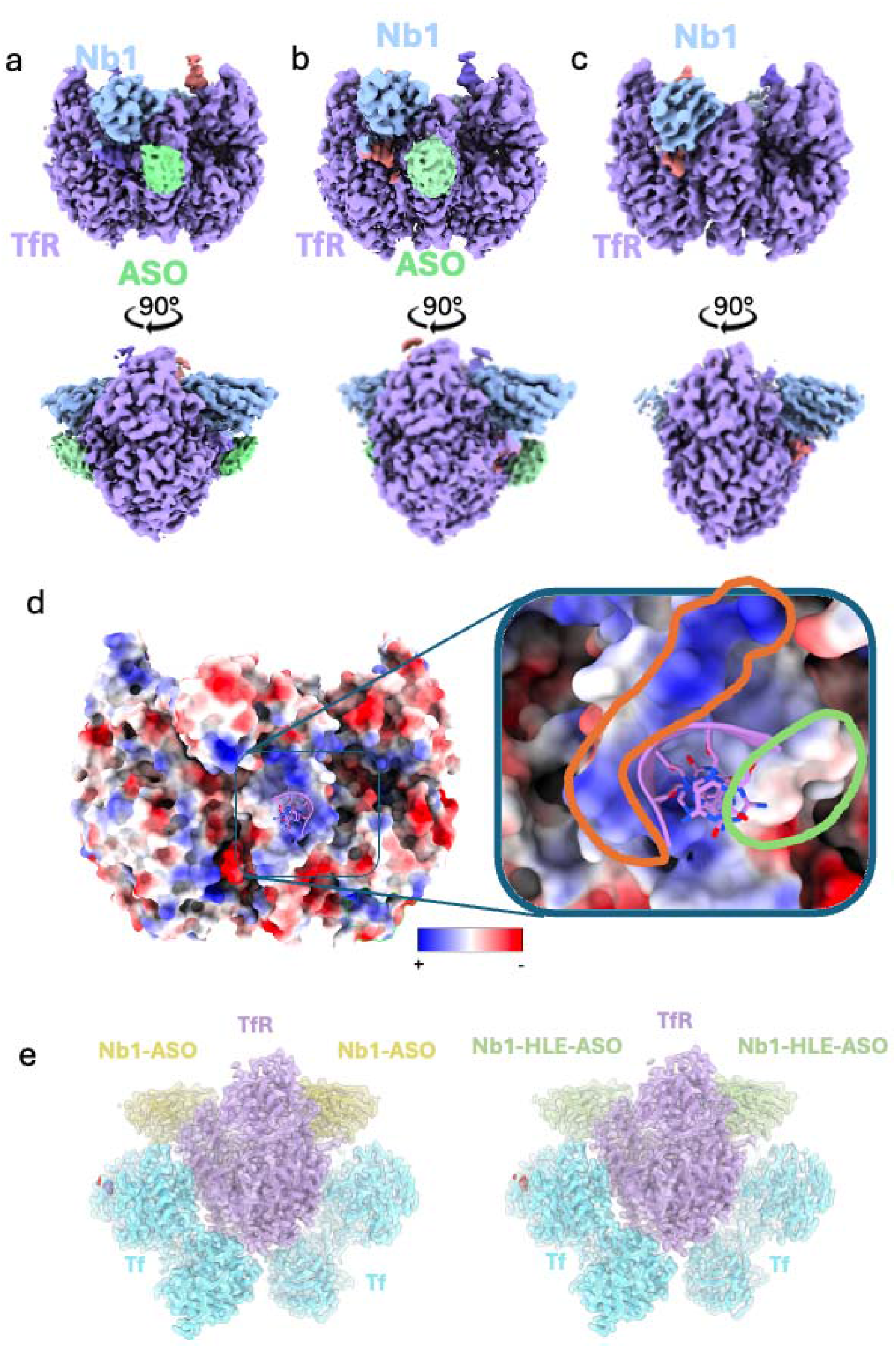
Structural analysis of anti-TfR1 NANOBODY® molecule Nb1 in complex with ASO reveals additional density corresponding to oligonucleotide binding. (a-b) Unsharpened cryo-EM density of the (a) TfR1+Nb1-Malat ASO and (b) TfR1+Nb1-Malat ASO complex showing extra density (highlighted in light green) that does not correspond to either the TfR1 receptor or the NANOBODY ® molecule. The density is shown in two orientations (90° rotation along the y axes). ChimeraX contour level used: 0.1. (c) Unsharpened cryo-EM density of the TfR-Nb1-siHprt complex showing no extra density can been seen in the transferrin-binding domain. The density is shown in two orientations (90° rotation along the y axis). Level used for visualization 0.074. (d) Coulombic rendering of the TfR1 surface with red showing negatively charged amino acids and positive ones in blue. The inset shows a magnified view of the binding interface where the positively charged surface (circled in orange) can accommodate the negatively charged phosphate backbone of the ASO, while hydrophobic region (circled in green) provides favourable interactions for the nucleotide bases. DNA was computationally docked using Boltz-2 and manually refined to optimize fit within the observed density. (e) Transparent cryo-EM density with fitted Ribbon diagram of the TfR-Tf-Nb1-ASO (left) and the TfR-Tf-Nb1-HLE-ASO (right) complexes at saturating concentrations of both Nb1 and Tf. Both Nb1 and Tf are clearly visible in the density and were built in the model.

Further analysis of the Nb1-ASO-hTfR1 complex using computational docking with Boltz2 ^34^, followed by manual refinement, positioned a ssDNA molecule with the ASO sequence within this density, providing a structural model for ASO binding^34^. Coulombic surface analysis of the TfR1 helical domain revealed an amphipathic region characterized by positively charged patches that could accommodate the negatively charged phosphate backbone of the ASO, while adjacent hydrophobic regions provide favorable interactions for the nucleotide bases (Figure 5d). This structural arrangement suggests a mechanism by which ASO payloads could directly contribute to TfR1 binding and indicates why the in vitro binding data showed a tighter affinity.

Importantly, even with the ASO payload’s ability to directly interact with TfR1, saturated complexes of TfR1 interacting with both Nb1-ASO conjugates and holo-Tf were still observed (Figure 5e). This indicates that the ASO interaction with TfR1 does not prevent holo-Tf from binding and vice versa for both Nb1-ASO conjugates and Nb1-HLE-ASO conjugates. An additional BLI binding experiment was also performed with each conjugate to further confirm that Nb1-HLE-ASO and Nb1-HLE-siRNA conjugates could still bind hTfR1 when saturating levels of holo-Tf were present, and conversely, that holo-Tf could still bind hTfR1 after saturating the hTfR1 with Nb1-HLE-ASO or Nb1-HLE-siRNA conjugates (Supplementary Figure 5).

### anti-TfR1 NANOBODY® molecule mediated delivery of siRNA leads to KD in skeletal and cardiac muscle

With the aim of evaluating whether anti-TfR1 NANOBODY® molecules can deliver siRNA into tissues that are traditionally considered hard to target, such as the skeletal and cardiac muscles, anti-TfR1 NANOBODY® molecules or negative control NANOBODY® molecules were conjugated to an siRNA targeting Hprt (siHprt). 200 nmol/kg of conjugated siHprt were dosed IV into hTfR1-KI mice, and tissues were collected 48h, 72h or 1 week after the single dose. Hprt knock down (KD) qPCR quantification revealed that ∼50% of target KD is achieved in quadriceps (Figure 6a) and gastrocnemius (Figure 6b) as early as 72h post IV dosing. The extent of target KD was lower in the heart (∼20%, Figure 6c).

**Figure 6.**
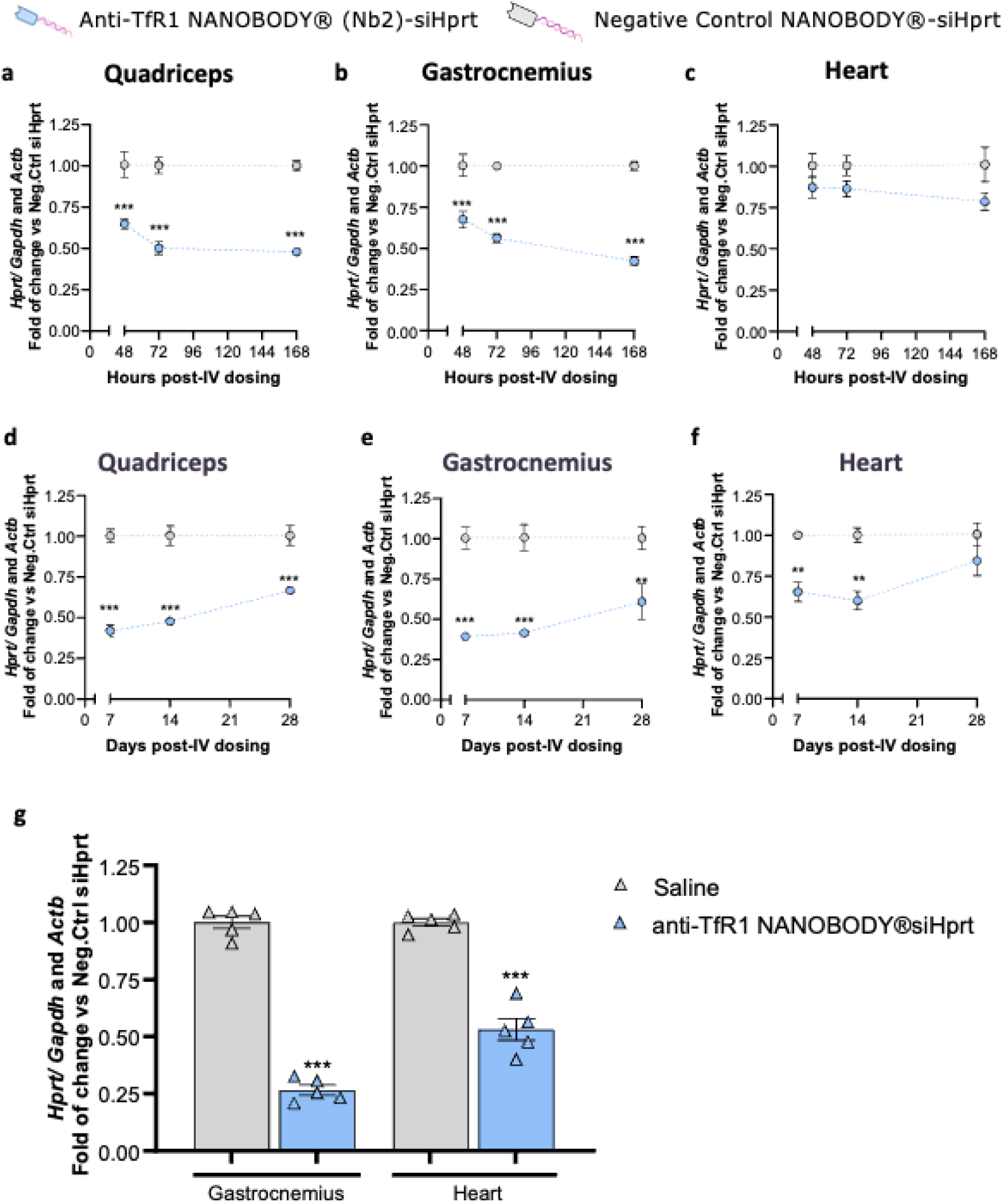
Anti-TfR1 NANOBODY® (Nb2)-siHprt leads to target KD in the muscle. (**a-c**) hTfR-KI mice were single IV dosed with 200nmol/kg NANOBODY®-siHprt conjugates, and (**a**) quadriceps, (**b**) gastrocnemius and (**c**) heart were collected 48h, 72h and 1 week post dosing. (**d-f**) hTfR-KI mice were single IV dosed with 200nmol/kg NANOBODY®-siHprt conjugates, and (**d**) quadriceps, (**e**) gastrocnemius and (f) heart were collected 1-, 2-and 4-weeks post dosing. (**g**) Repeated dosing of anti-TfR NANOBODY®-siHprt conjugates led to enhanced target KD in skeletal muscle and heart, as compared to single dose. hTfR-KI mice were dosed with 200nmol/kg anti-TfR NANOBODY®-siHprt conjugates once per week for four weeks and taken down one week after the fourth dose. Data is represented as Hprt levels fold change (2^-ΔΔCt^) vs negative control NANOBODY®-siHprt conjugate or saline. ΔCt of Hprt with Gapdh and Actb is calculated, followed by the geometric mean of these two ΔCt values. Mean +/- SEM represented, n=3 or n=5. Statistics: (a-f) 2 Way ANOVA, multiple comparisons test and (g) unpaired T test versus saline. ***= p<0.001

To better understand the longer-term duration of the target KD after a single siRNA dose upon conjugation to an anti-TfR1 NANOBODY®, hTfR1-KI mice were IV dosed with 200nmol/kg of conjugated siHprt, and mice were taken down after 1, 2 or 4 weeks. Quadriceps (Figure 6d) and gastrocnemius (Figure 6e) followed similar kinetics, in which maximal KD was observed between 1 and 2 weeks after dosing. The extent of the KD decreased by 4 weeks to ∼30% as compared to a negative control NANOBODY®-siHprt conjugate. Similar kinetics were determined in the heart (Figure 6f), with greatest KD of 30% between 1 and 2 weeks, while no KD was detectable at 4 weeks. Indicated by this dataset, it is worth noting that both target selection and tissue type can have critical impact on KD durability and magnitude.

Overall, these results demonstrate that a monovalent, non-half-life extended, anti-TfR1 NANOBODY® can deliver an oligonucleotide to the skeletal muscle, reaching a maximum KD effect up to 2 weeks after a single dose. However, to achieve a greater degree of target KD in organs impacted in various neuromuscular indications, a repeated dosing regimen with an anti-TfR1 NANOBODY® -siHprt conjugate was tested. For this, hTfR1-KI mice were dosed with 200nmol/kg of conjugated siRNA once a week for four weeks, and mice were taken down one week after the last dose. As shown in Figure 6g, repeated dosing of the conjugates increased the KD achieved in gastrocnemius (shown as a representative skeletal muscle) to 75%, while a 50% KD was determined in the heart. This % KD was greater than the efficacy shown after a single dose in Figures 6a-f, indicating a beneficial impact of a repeated dosing regimen.

### Half-life extension (HLE) is required for extended PK and enhanced brain exposure

With the goal of evaluating whether an extension in anti-TfR1 NANOBODY®’s half-life has an impact in brain exposure, anti-TfR1 NANOBODY® molecules with or without HLE, alongside an anti-TfR1 bispecific Ab (against TfR1 and an irrelevant target), were radiolabeled with ^125^I and IV dosed at equivalent molar doses into hTfR1-KI mice. For HLE purposes, a human serum albumin (HSA) binding NANOBODY® and an Fc were fused to an anti-TfR1 NANOBODY®.

Data in Figure 7a shows that while the plasma clearance of the non-HLE anti-TfR1 NANOBODY® is very fast, both types of HLE anti-TfR1 NANOBODY® formats (anti-HSA NANOBODY® or Fc) display prolonged plasma PK profiles, similar to the anti-TfR1 bispecific Ab control. Similarly, the brain exposure of the non-HLE anti-TfR1 NANOBODY® is lower than both HLE anti-TfR1 NANOBODY® formats, which reach similar concentrations and clearance kinetics as the anti-TfR1 bispecific Ab control (Figure 7b).

**Figure 7.**
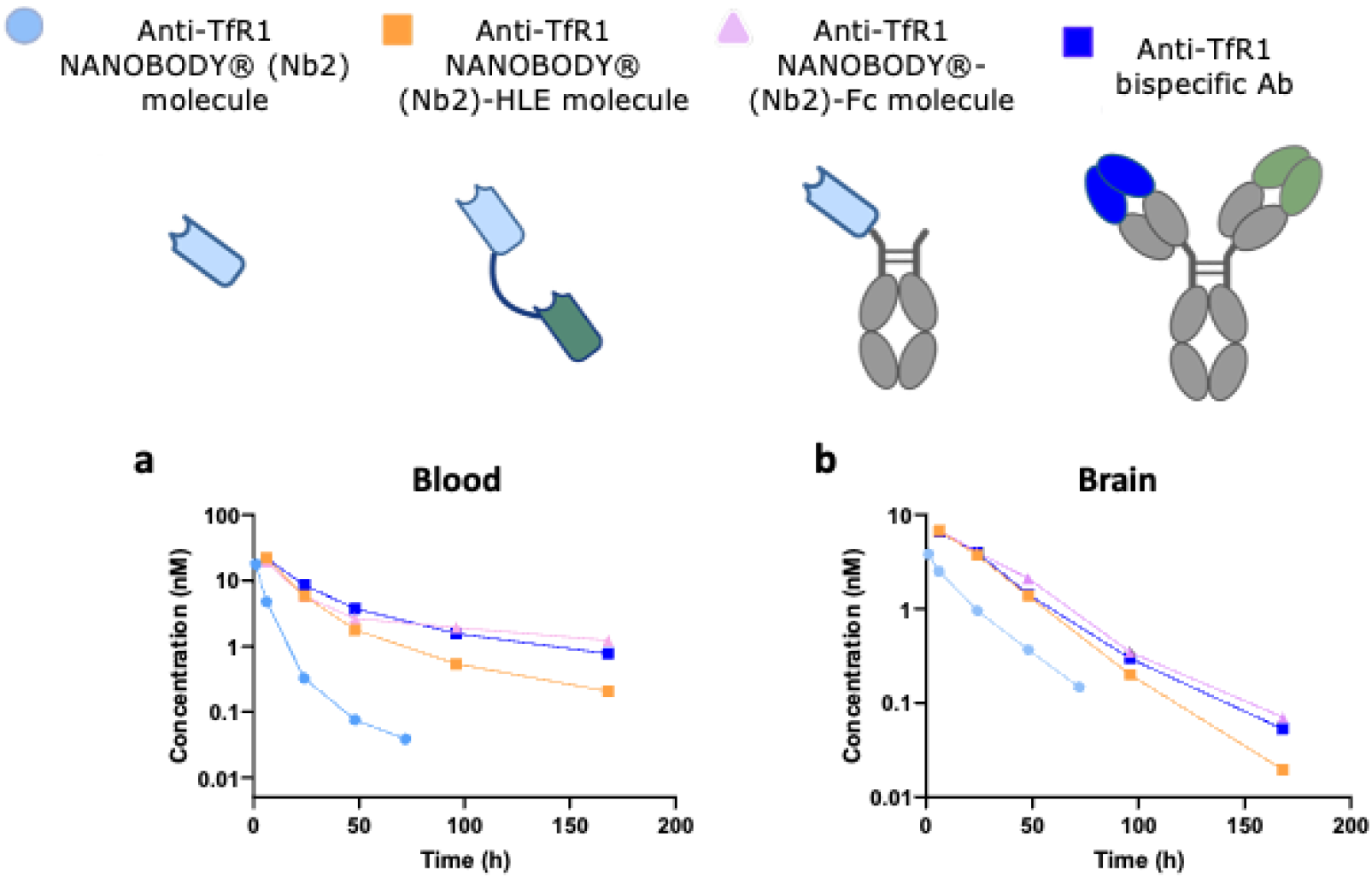
Plasma PK and brain exposure are enhanced with half-life extended anti-TfR1 NANOBODY® (Nb2) molecules. anti-TfR1 NANOBODY® (Nb2) molecules with or without half-life extension (HLE), alongside anti-TfR1 bispecific Ab (against TfR1 and an irrelevant target), were radiolabeled with ^125^I and IV dosed at equivalent molar doses (7 nmol/kg) into hTfR-KI mice. For HLE purposes, a human serum albumin (HSA) binding NANOBODY® molecule and an Fc were tested. Gamma count was quantified to determine tissue concentration of each anti-TfR1 construct. Mean +/- SEM represented, n=3.

This data indicates a beneficial effect of half-life extension for prolonging plasma PK and enhancing brain penetration of the anti-TfR1 NANOBODY molecules.

### Both anti-TfR1 NANOBODY®-HLE and anti-TfR1 NANOBODY®-HLE-Malat1 ASO conjugates successfully transcytose the BBB and show internalization in neurons

In order to evaluate whether ASO conjugation to anti-TfR1 NANOBODY® has an impact in BBB transcytosis and brain parenchymal cell internalization, both anti-TfR1 NANOBODY®-HLE and anti-TfR1 NANOBODY®-HLE-Malat1 ASO conjugates were administered as a single IV dose into hTfR1-KI mice and brains were collected post-perfusion at 24h. Immunofluorescence co-staining for VHH and NeuN revealed that both ASO conjugated and non-conjugated anti-TfR1 NANOBODY®-HLE molecules get internalized into neurons, which ensures the potential for ASO delivery into these brain parenchymal cells (Figure 8). Additional experiments with alternative NANOBODY® building blocks have revealed faster plasma half-lives for ASO conjugates when compared to unconjugated molecules. However, comparable brain exposure was still achieved for anti-TfR1 NANOBODY®-ASO conjugates (data not shown).

**Figure 8.**
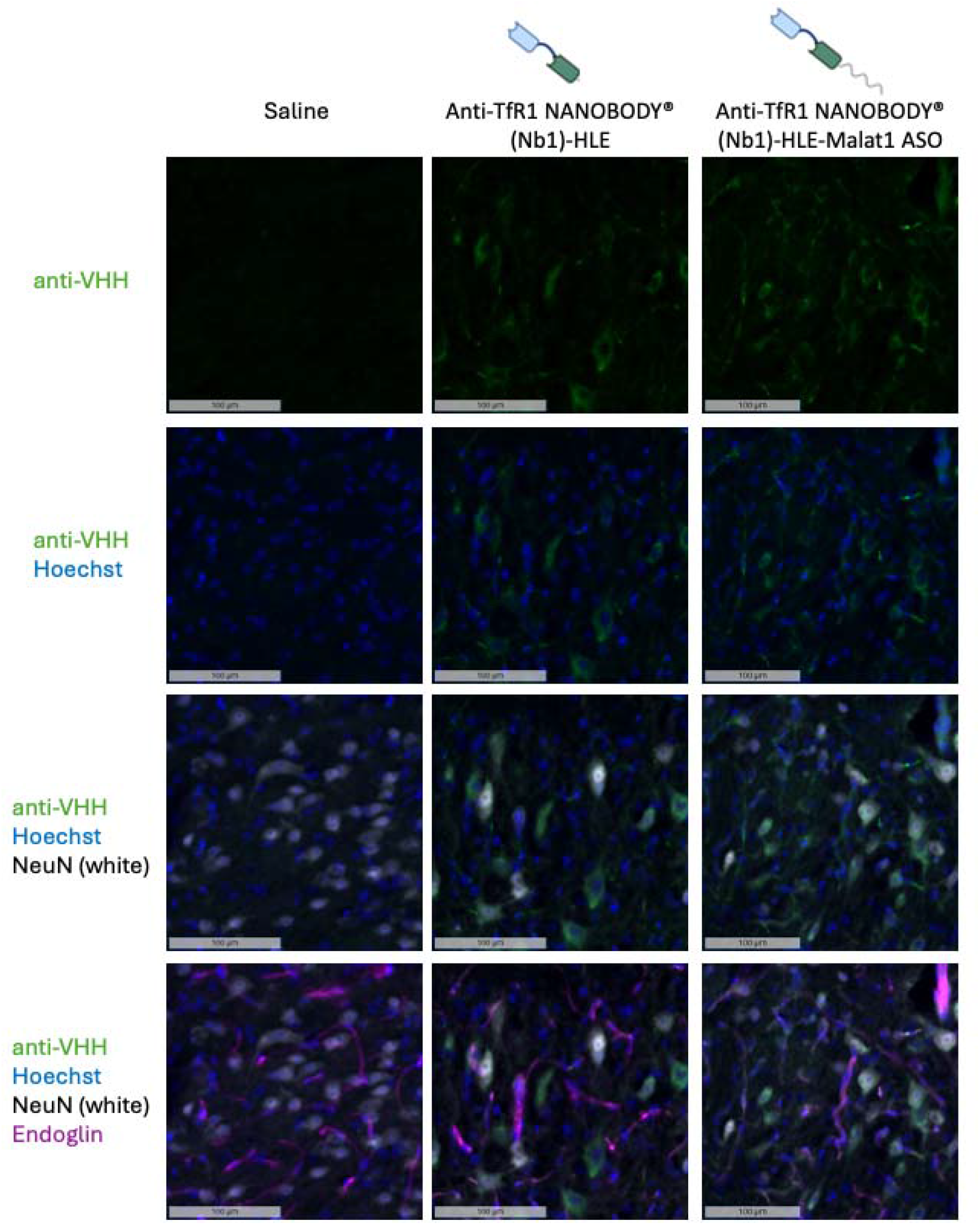
Anti-TfR1 NANOBODY® (Nb1) molecules both with and without Malat1 ASO conjugation show neuronal staining in hTfR-KI mice. hTfR-KI mice were single IV dosed with 200nmol/kg half-life extended (HLE) anti-TfR NANOBODY® (Nb1) constructs with and without ASO conjugation. At 24 h post-dose, brain hemispheres were collected, fixed, embedded, and stained. The saline treatment controls were then compared to the protein and conjugate treatment groups.

### anti-TfR1 NANOBODY®-HLE molecules deliver an ASO payload to the muscles, sciatic nerve and CNS, leading to target KD

Once increased brain exposure of anti-TfR1 NANOBODY®-HLE molecules was confirmed, their ability to deliver functional payload to tissues of interest (e.g. skeletal muscle, heart, sciatic nerve and brain) was evaluated. For this, anti-TfR1 NANOBODY® molecules with or without an HSA binding NANOBODY® for HLE, alongside a non-binding control NANOBODY® molecule, were conjugated to a Malat1 targeting Gapmer ASO (Figure 9). These conjugates were repeatedly dosed into hTfR1-KI mice, and after necropsy, Malat1 expression in gastrocnemius, quadriceps, heart, sciatic nerve and brain cortex was determined by qPCR. Malat1 ASO treated mice as well as mice treated with a non-targeted negative control NANOBODY® conjugate, did not display significant Malat1 KD in any tissue as compared to saline dosed mice (Figure 9b-f). When comparing the Malat1 KD levels achieved with anti-TfR1 NANOBODY® conjugates with or without HLE, in the quadriceps (Figure 9b) and gastrocnemius (Figure 9c) HLE and non-HLE formats lead to ∼75% target KD 72 h post last dose. snRNA-Seq from gastrocnemius tissue revealed target KD across all annotated cell types (Supplementary Figure 6a-b). Interestingly, presence of the HLE moiety led to a greater Malat1 KD in heart (Figure 9d), sciatic nerve (Figure 9e) and brain (Figure 9f), as compared to the non-HLE anti-TfR1 NANOBODY® molecule.

**Figure 9.**
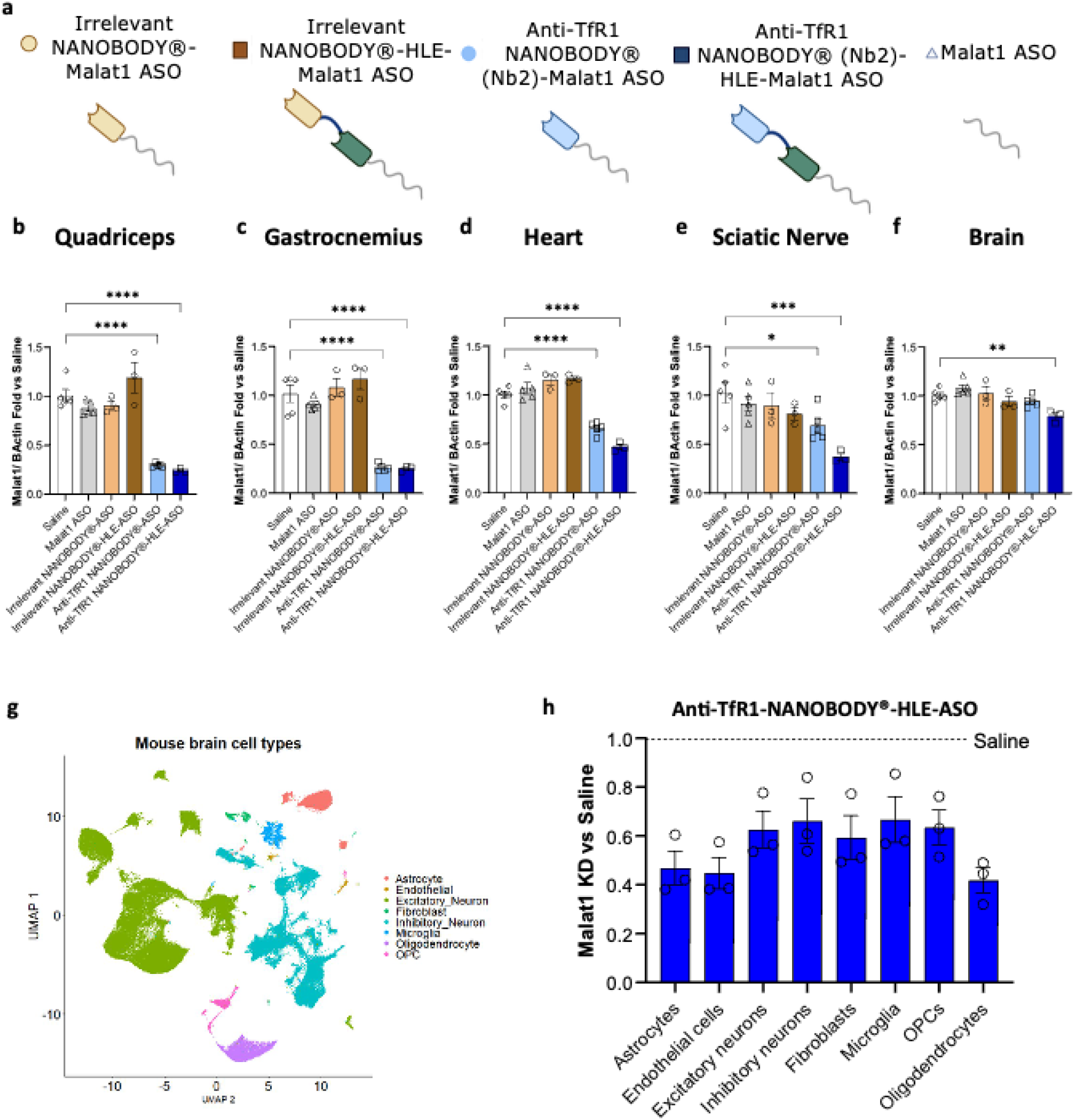
Half-life extended (HLE) anti-TfR1 NANOBODY® (Nb2) constructs deliver a Malat1- targting ASO payload to skeletal muscle, heart, sciatic nerve and CNS, resulting in target knockdown. Study details: hTfR-KI mice were IV dosed four times with 400 nmol/kg per dose over the course of 2 weeks with tissue collection 3 days post last dose. (a) Schematic representation of the molecules administered to hTfR-KI mice. (b-e) qRT-PCR analysis demonstrated statistically significant reduction of Malat1 expression across tissues in mice treated with anti-TfR1 NANOBODY® ± HLE Malat1 ASO constructs (n = 3-5). (f) Statistically significant reduction of Malat1 expression in brain tissue was achieved only with the anti-TfR1 NANOBODY® + HLE Malat1 ASO construct. Data were normalized to Actb and presented as fold change relative to saline. Statistical analysis was performed using one-way ANOVA followed by Dunnett’s multiple comparisions test versus saline. (g) UMAP representation of cells clusters identified by single-nucleus RNA sequencing from brain tissue. (h) Pseudo-bulk analysis from single-nucleus RNA sequencing data shows reduction of Malat1 expression across all annotated brain cell types. Mean +/- SEM represented, n=3.

Focusing on Malat1 KD in the brain, we further explored the specific brain cell types that the anti-TfR1 NANOBODY®-HLE molecules deliver the ASO to. For this, snRNA-Seq was performed (Figure 9g) followed by pseudo-bulk analysis; data indicated that all brain cell types within the brain can be targeted by anti-TfR1 NANOBODY®-HLE molecules, leading to a 30-60% KD of Malat1 across all cell types (Figure 9h).

Altogether, we have identified pH-dependent anti-mouse/ cyno/ human TfR1-targeting NANOBODY® molecules, that are suitable to use as shuttles to deliver therapeutic oligonucleotides into muscle and the CNS for the treatment of neuromuscular indications.

## Discussion

We report the discovery and characterization of pH-dependent, triple species cross-reactive anti-TfR1 NANOBODY® molecules that enable efficient oligonucleotide delivery to skeletal muscle and the CNS. These molecules exhibit several key features that make them promising therapeutic shuttles: (1) human, cyno, and mouse TfR1 cross-reactivity to facilitate preclinical-to-clinical translation; (2) pH-dependent binding to promote both surface engagement and endosomal release; (3) non-competitive binding with transferrin to preserve iron homeostasis; and (4) efficient payload delivery to traditionally hard-to-target tissues.

### Structural basis for pH-dependent binding and therapeutic implications

The three NANOBODY® molecules discussed in this work (Nb1-Nb3) recognize overlapping epitopes at the TfR1 apical helical domain, with contacts extending to the protease-like domain of the adjacent subunit. Comparative structural analysis at pH 7.4 and pH 6.0 uncovered the molecular mechanism underlying pH-dependent binding: a histidine-mediated conformational switch involving His318 and His475. At physiological pH, the His318-containing loop adopts a compact conformation, leaving the NANOBODY® epitope accessible. Upon endosomal acidification, protonation of His318 triggers a displacement of this loop, sterically occluding the epitope and promoting NANOBODY® molecule dissociation.

The pH-dependent behavior we observe is dictated by the epitope location rather than the NANOBODY® paratope design, as all three clones exhibit similar pH sensitivity despite subtle differences in their interaction patterns. This represents a distinct mechanism from engineered pH-sensitivity approaches and suggests that natural pH-responsive epitopes on TfR1 can be exploited for therapeutic advantage. The His318 loop functions as a natural pH-responsive “lid” that likely plays a regulatory role in TfR1’s endocytic cycle, potentially coordinating iron release from transferrin or facilitating receptor recycling.^16^

This pH-dependent mechanism offers distinct advantages for CNS delivery by addressing the “binding site barrier” phenomenon, where high-affinity molecules become trapped within brain vasculature rather than achieving parenchymal penetration^11–13,35^ Strong binding at neutral pH (cell surface) ensures efficient receptor engagement and internalization, while weakened binding at acidic pH in endosomes, facilitates cargo release and deeper tissue penetration. Our findings align with recent reports demonstrating the importance of pH-sensitive binding for BBB transcytosis.^14,15^

### Comparison with established TfR1-targeting platforms

Our anti-TfR1 NANOBODY® molecules complement the growing arsenal of single-domain antibodies targeting TfR1 for CNS delivery. The field has established that single-domain antibodies (e.g. VHHs or VNARs) offer significant advantages as “molecular shuttles” due to their small size, high stability, and ability to access cryptic epitopes often inaccessible to conventional monoclonal antibodies.^36–38^ In a similar example, engineered VHH molecules demonstrated successful TfR1-dependent transcytosis across the BBB when fused to therapeutic payloads, exhibiting favorable pharmacokinetics and minimal hematotoxicity.^39,40^ A recent study also reported novel VHHs targeting a unique TfR1 epitope with cross-species reactivity and significantly improved brain uptake in mouse models.^41,42^ Our anti-TfR1 NANOBODY® molecules target the same general TfR1 dimer interface, and, consistent with their report, we also observe triple species cross-reactivity and pH-dependent TfR1 binding.

Studies have established that the efficacy of TfR1-mediated delivery is highly dependent on molecular architecture. ^6,11–13^ While high affinity is often desired, intermediate affinity and rapid dissociation rates have been associated with more efficient BBB transcytosis, as they prevent molecules from being trapped within brain vasculature. ^6,35^ Our molecules, with single-digit nanomolar to sub-nanomolar affinity at pH 7.4 and up to 250-fold reduced binding at pH 6.0, fall within the optimal range predicted by these models. The combination of intermediate effective affinity, pH-sensitivity, and monovalent binding format positions our platform favorably for CNS applications^6^. However, our structural characterization provides unprecedented mechanistic insight into the pH-dependent conformational changes that drive epitope accessibility.

### Epitope selection and transferrin competition

Epitope binning confirmed that Nb1-Nb3 compete for the same binding site but do not interfere with transferrin binding. This non-competitive binding profile is critical for therapeutic safety, as disruption of iron homeostasis could lead to systemic toxicity. The consistent finding across multiple independent TfR1-targeting platforms, including the NewroBus VHHs, cross-reactive VHHs, and shark-derived VNARs, that epitopes at the TfR1 dimer interface do not interfere with iron homeostasis suggests that this region represents a privileged site for therapeutic targeting.^36,37,39,42^. The structural conservation of this region across species (human, cyno, mouse) further supports its suitability for cross-reactive binder development, addressing the critical need for platforms that enable seamless translation from preclinical models to clinical applications.^39,40,42^

### ASO-TfR1 interactions enhance binding affinity but do not prevent Tf binding

An unexpected finding from our structural studies was an extra density in TfR1–NANOBODY®– ASO complexes, resulting from the direct interaction between single-stranded ASO payloads and the TfR1 helical domain. This additional electron density in Nb1-ASO-TfR1 complexes was absent in Nb1-siRNA-TfR1 complexes, indicating that only single-stranded oligonucleotides engage the receptor. This direct ASO-TfR1 interaction explains the ∼10-fold tighter binding affinity observed for ASO conjugates compared to unconjugated NANOBODY® molecules or siRNA conjugates.

To our knowledge, this is the first structural demonstration of direct oligonucleotide-TfR1 interactions in the context of antibody-oligonucleotide conjugates (AOCs). The design of TfR1-targeting oligonucleotide conjugates requires careful optimization of binding affinity and valency to ensure efficient parenchymal accumulation, as demonstrated by successful transport of antisense oligonucleotides across the mammalian blood-brain barrier.^9^ Our findings suggest that payload architecture: single-stranded vs. double-stranded, backbone modifications, sequence composition, may significantly impact delivery efficiency through direct receptor interactions, adding an additional layer of complexity to conjugate design. The phosphorothioate backbone modifications in our ASO, incorporated to enhance metabolic stability, likely strengthen these TfR1 interactions, consistent with previous reports showing increased protein binding by PS-modified oligonucleotides^43^. Importantly, ASO binding does not prevent transferrin engagement, preserving physiological iron transport while still exhibiting efficacy in skeletal muscle and CNS.

Recent studies have demonstrated successful delivery of antisense oligonucleotides to the CNS using TfR1-targeting constructs, but the molecular basis for how oligonucleotide payloads influence TfR1 binding has remained unclear.^9^ Our structural data provide the first mechanistic explanation for these observations.

### Pharmacokinetic optimization enables CNS delivery across multiple cell types

Our biodistribution studies demonstrate that the molecular format critically determines tissue exposure and functional payload delivery, consistent with established principles^11–13^. Non-half-life-extended anti-TfR1 NANOBODY®-siRNA conjugates achieved rapid and sustained knockdown in skeletal muscle (up to 60% at 2 weeks and 35-40% at 4 weeks post-single dose), with repeated dosing enhancing efficacy to 75% in gastrocnemius and 50% in heart. However, brain exposure of these short-circulating formats was limited, highlighting the need to explore half-life extension opportunities.

Incorporation of half-life extension, either through fusion to an anti-human serum albumin NANOBODY® or an Fc domain, dramatically improved both plasma pharmacokinetics and brain penetration. HLE anti-TfR1 NANOBODY®-ASO conjugates enabled significant target knockdown in heart, sciatic nerve, and brain cortex upon repeated dosing, with single-nucleus RNA sequencing revealing 30-60% Malat1 knockdown across all major brain cell types. Immunofluorescence confirmed neuronal internalization of both conjugated and unconjugated HLE formats, demonstrating successful BBB transcytosis and parenchymal delivery. This broad cell-type coverage addresses a key limitation of many CNS delivery platforms that show restricted cellular distribution.

These findings align with recent advances demonstrating that TfR1-targeting NANOBODY® molecules can serve as modular platforms for delivering various therapeutic modalities, including biologics, enzymes, and oligonucleotides.^6,9^ Anti-TfR1 NANOBODY® molecules have also been utilized as imaging tools, such as PET radioligands to track disease-relevant proteins or cells in vivo, including development of congenic mice for imaging transplants by positron emission tomography using anti-transferrin receptor NANOBODY® molecules and studies of passive versus receptor-mediated brain delivery, further demonstrating the versatility of this platform across therapeutic and diagnostic applications.^44,45^

The ability to achieve functional knockdown across diverse CNS cell types, without intrathecal administration, represents a significant advance for oligonucleotide-based neurotherapeutics. Previous studies have primarily focused on BBB penetration and total brain exposure, but cell-type-specific delivery and functional activity have been less well characterized.^6,9^ Our single-nucleus RNA sequencing approach provides definitive evidence that TfR1-mediated delivery can reach therapeutically relevant cell populations throughout the brain parenchyma.

### Conclusion

In this work, we have reported the discovery and validation of anti-TfR1 NANOBODY® molecules (Nb1-Nb3) for delivery of oligonucleotide therapeutics to skeletal muscle and CNS tissues. These NANOBODY® molecules leverage triple species cross-reactivity as well as pH-dependent binding to TfR, as further explained through the structural datasets presented. While this work shows the promise of NANOBODY® technology and modality for oligonucleotide delivery, future work is required to further determine therapeutic translatability. This future work should focus on: (1) validating efficacy with a variety of oligo payloads in disease-relevant models of neuromuscular and neurodegenerative disorders, building on the clinical success of TfR1-targeted oligonucleotide delivery platforms ^9^; (2) optimizing dosing regimens guided by molecular architecture principles to balance efficacy, durability, and safety ^6,35^; (3) engineering NANOBODY® variants with tunable pH sensitivity for specific applications. This study adds to the growing field of evidence exhibiting VHHs and our NANOBODY® molecules as viable shuttles to deliver oligonucleotide therapeutics and provide improved options and impact for patients.

## Data Availability

The cryo-EM density maps (including independent half-maps, unsharpened maps, and sharpened maps) have been deposited in the Electron Microscopy Data Bank under accession code EMD-XXXXX. The atomic coordinates have been deposited in the Protein Data Bank under accession code XXXX.

The sequencing data generated in this study will be deposited in the NCBI Gene Expression Omnibus (GEO) and made publicly available upon publication.

## Acknowledgments

NANOBODY® is a registered trademark of Ablynx NV, an affiliate of Sanofi.

## Authors Contributions

E.A.H., K.M., C.R., A.M.S., S.C. and N.L. designed the study. K.M., C.N. and S.C. designed and supervised the workflow related to NANOBODY® identification and characterization, including immunization, phage display, primary screening, NANOBODY® expression/purification, MSD, and transferrin competition assays. E.A.H., A.I., F.M. and J.T.conducted biophysical characterization including BLI, epitope binning, and transferrin competition assays. C.C., F.M., and S.R. expressed TfR1 protein from different species. C.R. and G.H., conducted cryo-EM structure determination and structural data analysis. E.A.H., A.M.S., R.M.-G., F.M., S.R., J.T., S.Z. and A.I. conducted NANOBODY®–oligonucleotide conjugation and in vivo knockdown studies.

S.K. and L.A. conducted radiolabeled biodistribution and pharmacokinetic studies. E.A.H., A.M.S. and R.M.-G. coordinated brain parenchymal biodistribution and immunofluorescence imaging. T.R.H. and R.M.-G. conducted single-nucleus RNA sequencing and analysis. P.S., S.C. and N.L. supervised the research. E.A.H., K.M., C.R., A.M.S., T.R.H., S.K., G.H., C.C., F.M., L.A., C.N., S.C. and N.L. interpreted the data. E.A.H., K.M., C.R., A.M.S., C.S., and N.L. wrote the paper with input from the other authors.

## References

1. Thornton, C. A. et al. Antisense oligonucleotide targeting DMPK in patients with myotonic dystrophy type 1: a multicentre, randomised, dose-escalation, placebo-controlled, phase 1/2a trial. Lancet Neurol. 22, 218–228 (2023).

2. Christensen, J. et al. Metabolism Studies of Unformulated Internally [3H]-Labeled Short Interfering RNAs in Mice. Drug Metab. Dispos. 41, 1211–1219 (2013).

3. Godinho, B. M. D. C. et al. PK-modifying anchors significantly alter clearance kinetics, tissue distribution, and efficacy of therapeutics siRNAs. Mol. Ther. - Nucleic Acids 29, 116–132 (2022).

4. Baik, A. D. et al. Targeted delivery of acid alpha-glucosidase corrects skeletal muscle phenotypes in Pompe disease mice. bioRxiv 2020.04.22.051672 (2020) doi:10.1101/2020.04.22.051672.

5. George, K. et al. Novel transferrin receptor-mediated enzyme replacement therapy efficiently treats myogenic and neurogenic aspects of Pompe disease in mice. Mol. Ther. Methods Clin. Dev. 33, 101547 (2025).

6. Arguello, A. et al. Molecular architecture determines brain delivery of a transferrin receptor– targeted lysosomal enzyme. J. Exp. Med. 219, e20211057 (2022).

7. Gehrlein, A. et al. Targeting neuronal lysosomal dysfunction caused by β-glucocerebrosidase deficiency with an enzyme-based Brain Shuttle construct. (2022) doi:10.21203/rs.3.rs-1490073/v1.

8. Lengerich, B. van, et al. A TREM2-activating antibody with a blood–brain barrier transport vehicle enhances microglial metabolism in Alzheimer’s disease models. Nat. Neurosci. 26, 416– 429 (2023).

9. Barker, S. J. et al. Targeting the transferrin receptor to transport antisense oligonucleotides across the mammalian blood-brain barrier. Sci. Transl. Med. 16, eadi2245 (2024).

10. Weeden, T. et al. FORCE platform overcomes barriers of oligonucleotide delivery to muscle and corrects myotonic dystrophy features in preclinical models. Commun. Med. 5, 22 (2025).

11. Niewoehner, J. et al. Increased Brain Penetration and Potency of a Therapeutic Antibody Using a Monovalent Molecular Shuttle. Neuron 81, 49–60 (2014).

12. Rosa, A. de la, et al. Lowering the affinity of single-chain monovalent BBB shuttle scFc-scFv8D3 prolongs its half-life and increases brain concentration. Neurotherapeutics 22, e00492 (2025).

13. Yu, Y. J. et al. Therapeutic bispecific antibodies cross the blood-brain barrier in nonhuman primates. Sci. Transl. Med. 6, 261ra154 (2014).

14. Sade, H. et al. A Human Blood-Brain Barrier Transcytosis Assay Reveals Antibody Transcytosis Influenced by pH-Dependent Receptor Binding. PLoS ONE 9, e96340 (2014).

15. Esparza, T. J. et al. Enhanced in vivo blood brain barrier transcytosis of macromolecular cargo using an engineered pH-sensitive mouse transferrin receptor binding nanobody. Fluids Barriers CNS 20, 64 (2023).

16. Steere, A. N. et al. Structure-Based Mutagenesis Reveals Critical Residues in the Transferrin Receptor Participating in the Mechanism of pH-Induced Release of Iron from Human Serum Transferrin. Biochemistry 51, 2113–2121 (2012).

17. Groeve, M. D., Laukens, B. & Schotte, P. Optimizing expression of Nanobody® molecules in Pichia pastoris through co-expression of auxiliary proteins under methanol and methanol-free conditions. Microb. Cell Factories 22, 135 (2023).

18. Punjani, A., Rubinstein, J. L., Fleet, D. J. & Brubaker, M. A. cryoSPARC: algorithms for rapid unsupervised cryo-EM structure determination. Nat. Methods 14, 290–296 (2017).

19. Punjani, A., Zhang, H. & Fleet, D. J. Non-uniform refinement: adaptive regularization improves single-particle cryo-EM reconstruction. Nat. Methods 17, 1214–1221 (2020).

20. Scheres, S. H. W. RELION: Implementation of a Bayesian approach to cryo-EM structure determination. J. Struct. Biol. 180, 519–530 (2012).

21. Kimanius, D., Forsberg, B. O., Scheres, S. H. & Lindahl, E. Accelerated cryo-EM structure determination with parallelisation using GPUs in RELION-2. eLife 5, e18722 (2016).

22. Chen, S. et al. High-resolution noise substitution to measure overfitting and validate resolution in 3D structure determination by single particle electron cryomicroscopy. Ultramicroscopy 135, 24–35 (2013).

23. Terwilliger, T. C., Adams, P. D., Afonine, P. V. & Sobolev, O. V. A fully automatic method yielding initial models from high-resolution cryo-electron microscopy maps. Nat. Methods 15, 905–908 (2018).

24. Sanchez-Garcia, R. et al. DeepEMhancer: a deep learning solution for cryo-EM volume post-processing. Commun. Biol. 4, 874 (2021).

25. Jumper, J. et al. Highly accurate protein structure prediction with AlphaFold. Nature 596, 583–589 (2021).

26. Emsley, P., Lohkamp, B., Scott, W. G. & Cowtan, K. Features and development of Coot. Acta Crystallogr. Sect. D: Biol. Crystallogr. 66, 486–501 (2010).

27. Croll, T. I. ISOLDE: a physically realistic environment for model building into low-resolution electron-density maps. Acta Crystallogr. Sect. D: Struct. Biol. 74, 519–530 (2018).

28. Afonine, P. V. et al. Real-space refinement in PHENIX for cryo-EM and crystallography. Acta Crystallogr. Sect. D: Struct. Biol. 74, 531–544 (2018).

29. Goddard, T. D. et al. UCSF ChimeraX: Meeting modern challenges in visualization and analysis. Protein Sci. 27, 14–25 (2018).

30. Young, M. D. & Behjati, S. SoupX removes ambient RNA contamination from droplet-based single-cell RNA sequencing data. GigaScience 9, giaa151 (2020).

31. Yang, L. et al. Single-cell Mayo Map (scMayoMap): an easy-to-use tool for cell type annotation in single-cell RNA-sequencing data analysis. BMC Biol. 21, 223 (2023).

32. Dizicheh, Z. B., Chen, I.-L. & Koenig, P. VHH CDR-H3 conformation is determined by VH germline usage. Commun. Biol. 6, 864 (2023).

33. Sela, T. et al. Impact of ASO conjugation and receptor binding affinity on intracellular transport of mono- and bispecific TfR- and CD98-BrainshuttleTM variants. mAbs 18, 2691351 (2026).

34. Passaro, S., et al. Boltz-2: Towards Accurate and Efficient Binding Affinity Prediction. bioRxiv 2025.06.14.659707 (2025) doi:10.1101/2025.06.14.659707.

35. Thomsen, M. S. & Moos, T. A novel bispecific antibody able to pass the blood-brain barrier and therapeutically engage within the brain. Med 3, 815–817 (2022).

36. Clarke, E. et al. A Single Domain Shark Antibody Targeting the Transferrin Receptor 1 Delivers a TrkB Agonist Antibody to the Brain and Provides Full Neuroprotection in a Mouse Model of Parkinson’s Disease. Pharmaceutics 14, 1335 (2022).

37. Stocki, P. et al. Blood-brain barrier transport using a high affinity, brain-selective VNAR antibody targeting transferrin receptor 1. FASEB J. 35, e21172 (2021).

38. Zhou, L., et al. Self-Assembled Antibody-Oligonucleotide Conjugates for Targeted Delivery of Complementary Antisense Oligonucleotides. Angew. Chem. Int. Ed. 64, e202415272 (2025).

39. Yin, T. et al. The NewroBus platform: engineered humanized anti-TfR1 nanobodies for efficient brain delivery. Cell Commun. Signal. 24, 69 (2025).

40. Lemprière, S. Novel transport vehicle delivers biotherapeutics to the brain. Nat. Rev. Neurol. 16, 404–404 (2020).

41. Jacquot, G. et al. Harnessing TfR1 for Cross-Species Systemic Delivery of siRNAs to Deep Brain Regions Using Single-Domain Antibodies. bioRxiv 2026.05.20.726486 (2026) doi:10.64898/2026.05.20.726486.

42. David, M. et al. Novel single-domain antibodies targeting a unique transferrin receptor 1 epitope for cross-species delivery of drugs in the central nervous system. J. Control. Release 388, 114344 (2025).

43. Hyjek-Składanowska, M., et al. Origins of the Increased Affinity of Phosphorothioate-Modified Therapeutic Nucleic Acids for Proteins. J. Am. Chem. Soc. 142, 7456–7468 (2020).

44. Meier, S. R., Sehlin, D. & Syvänen, S. Passive and receptor mediated brain delivery of an anti-GFAP nanobody. Nucl. Med. Biol. 114, 128–134 (2022).

45. Balligand, T. et al. A pair of congenic mice for imaging of transplants by positron emission tomography using anti-transferrin receptor nanobodies. eLife 14, RP104302 (2025).

