## Supplementary Files for "pH-dependent anti-TfR1 NANOBODY® molecules deliver efficacious oligonucleotide payloads to muscle and CNS tissues"

### Slide 1
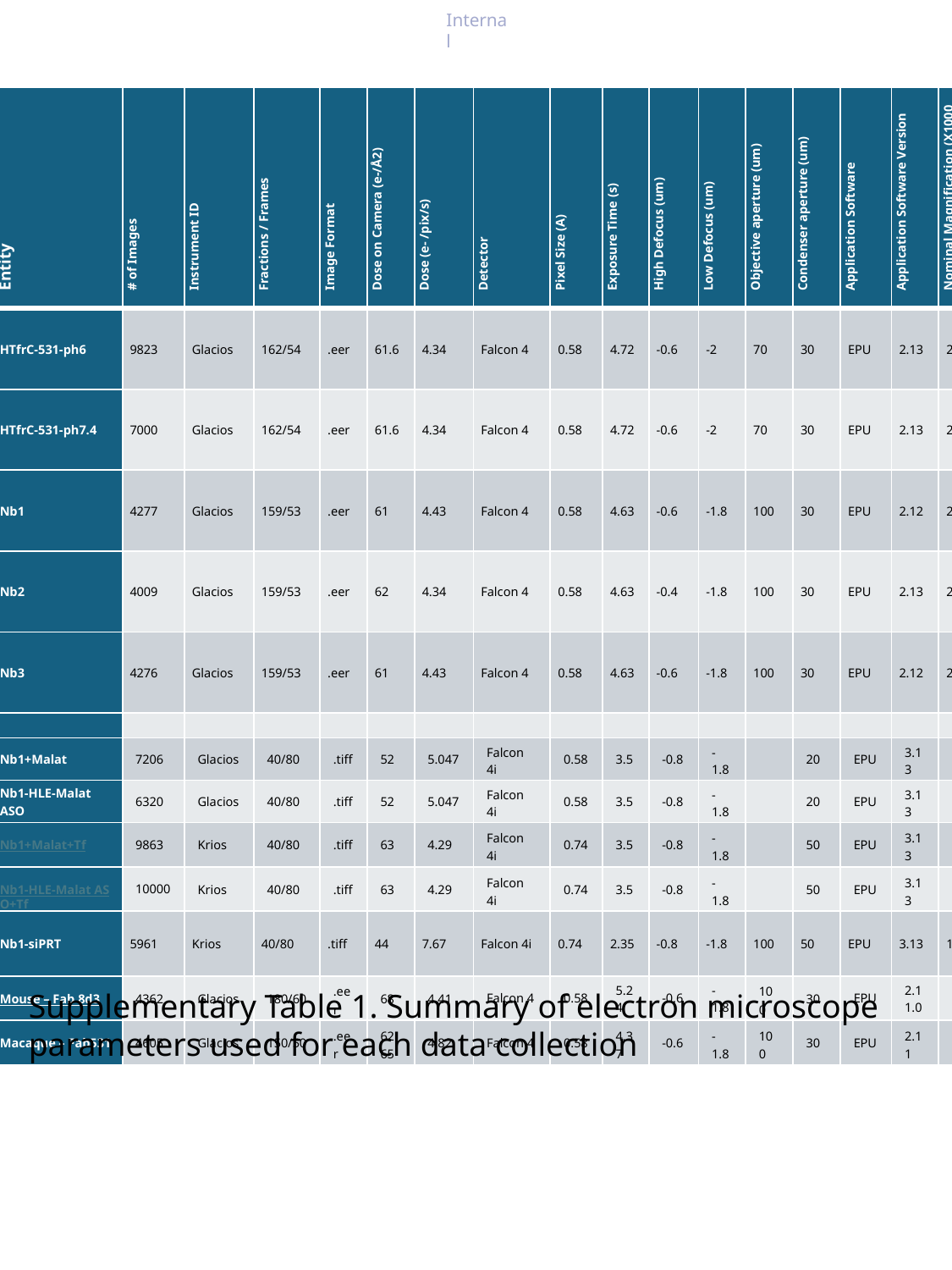

| Entity | # of Images | Instrument ID | Fractions / Frames | Image Format | Dose on Camera (e-/Å2) | Dose (e- /pix/s) | Detector | Pixel Size (A) | Exposure Time (s) | High Defocus (um) | Low Defocus (um) | Objective aperture (um) | Condenser aperture (um) | Application Software | Application Software Version | Nominal Magnification (X1000 X) |
| --- | --- | --- | --- | --- | --- | --- | --- | --- | --- | --- | --- | --- | --- | --- | --- | --- |
| HTfrC-531-ph6 | 9823 | Glacios | 162/54 | .eer | 61.6 | 4.34 | Falcon 4 | 0.58 | 4.72 | -0.6 | -2 | 70 | 30 | EPU | 2.13 | 240 |
| HTfrC-531-ph7.4 | 7000 | Glacios | 162/54 | .eer | 61.6 | 4.34 | Falcon 4 | 0.58 | 4.72 | -0.6 | -2 | 70 | 30 | EPU | 2.13 | 240 |
| Nb1 | 4277 | Glacios | 159/53 | .eer | 61 | 4.43 | Falcon 4 | 0.58 | 4.63 | -0.6 | -1.8 | 100 | 30 | EPU | 2.12 | 240 |
| Nb2 | 4009 | Glacios | 159/53 | .eer | 62 | 4.34 | Falcon 4 | 0.58 | 4.63 | -0.4 | -1.8 | 100 | 30 | EPU | 2.13 | 240 |
| Nb3 | 4276 | Glacios | 159/53 | .eer | 61 | 4.43 | Falcon 4 | 0.58 | 4.63 | -0.6 | -1.8 | 100 | 30 | EPU | 2.12 | 240 |
| Nb1+Malat | 7206 | Glacios | 40/80 | .tiff | 52 | 5.047 | Falcon 4i | 0.58 | 3.5 | -0.8 | -1.8 | | 20 | EPU | 3.13 | 205 |
| Nb1-HLE-Malat ASO | 6320 | Glacios | 40/80 | .tiff | 52 | 5.047 | Falcon 4i | 0.58 | 3.5 | -0.8 | -1.8 | | 20 | EPU | 3.13 | 205 |
| Nb1+Malat+Tf | 9863 | Krios | 40/80 | .tiff | 63 | 4.29 | Falcon 4i | 0.74 | 3.5 | -0.8 | -1.8 | | 50 | EPU | 3.13 | 165 |
| Nb1-HLE-Malat ASO+Tf | 10000 | Krios | 40/80 | .tiff | 63 | 4.29 | Falcon 4i | 0.74 | 3.5 | -0.8 | -1.8 | | 50 | EPU | 3.13 | 165 |
| Nb1-siPRT | 5961 | Krios | 40/80 | .tiff | 44 | 7.67 | Falcon 4i | 0.74 | 2.35 | -0.8 | -1.8 | 100 | 50 | EPU | 3.13 | 165 |
| Mouse – Fab 8d3 | 4362 | Glacios | 180/60 | .eer | 68 | 4.41 | Falcon 4 | 0.58 | 5.24 | -0.6 | -1.8 | 100 | 30 | EPU | 2.11.0 | 240 |
| Macaque – Fab531 | 4605 | Glacios | 150/50 | .eer | 62.65 | 4.82 | Falcon 4 | 0.58 | 4.37 | -0.6 | -1.8 | 100 | 30 | EPU | 2.11 | 240 |
Supplementary Table 1. Summary of electron microscope parameters used for each data collection

### Slide 2
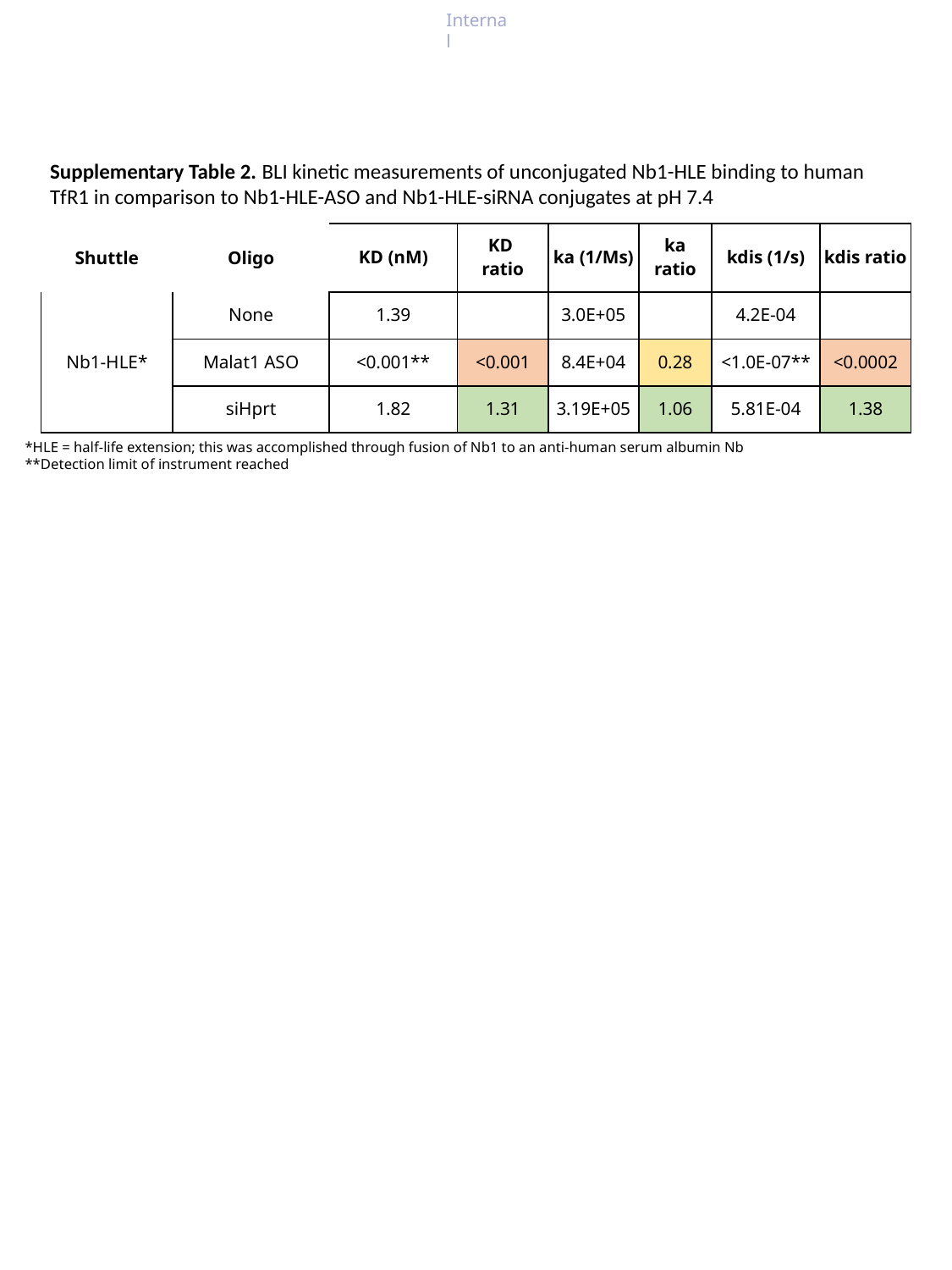

Supplementary Table 2. BLI kinetic measurements of unconjugated Nb1-HLE binding to human TfR1 in comparison to Nb1-HLE-ASO and Nb1-HLE-siRNA conjugates at pH 7.4
| Shuttle | Oligo | KD (nM) | KD ratio | ka (1/Ms) | ka ratio | kdis (1/s) | kdis ratio |
| --- | --- | --- | --- | --- | --- | --- | --- |
| Nb1-HLE\* | None | 1.39 | | 3.0E+05 | | 4.2E-04 | |
| | Malat1 ASO | <0.001\*\* | <0.001 | 8.4E+04 | 0.28 | <1.0E-07\*\* | <0.0002 |
| | siHprt | 1.82 | 1.31 | 3.19E+05 | 1.06 | 5.81E-04 | 1.38 |
*HLE = half-life extension; this was accomplished through fusion of Nb1 to an anti-human serum albumin Nb
**Detection limit of instrument reached

### Slide 3
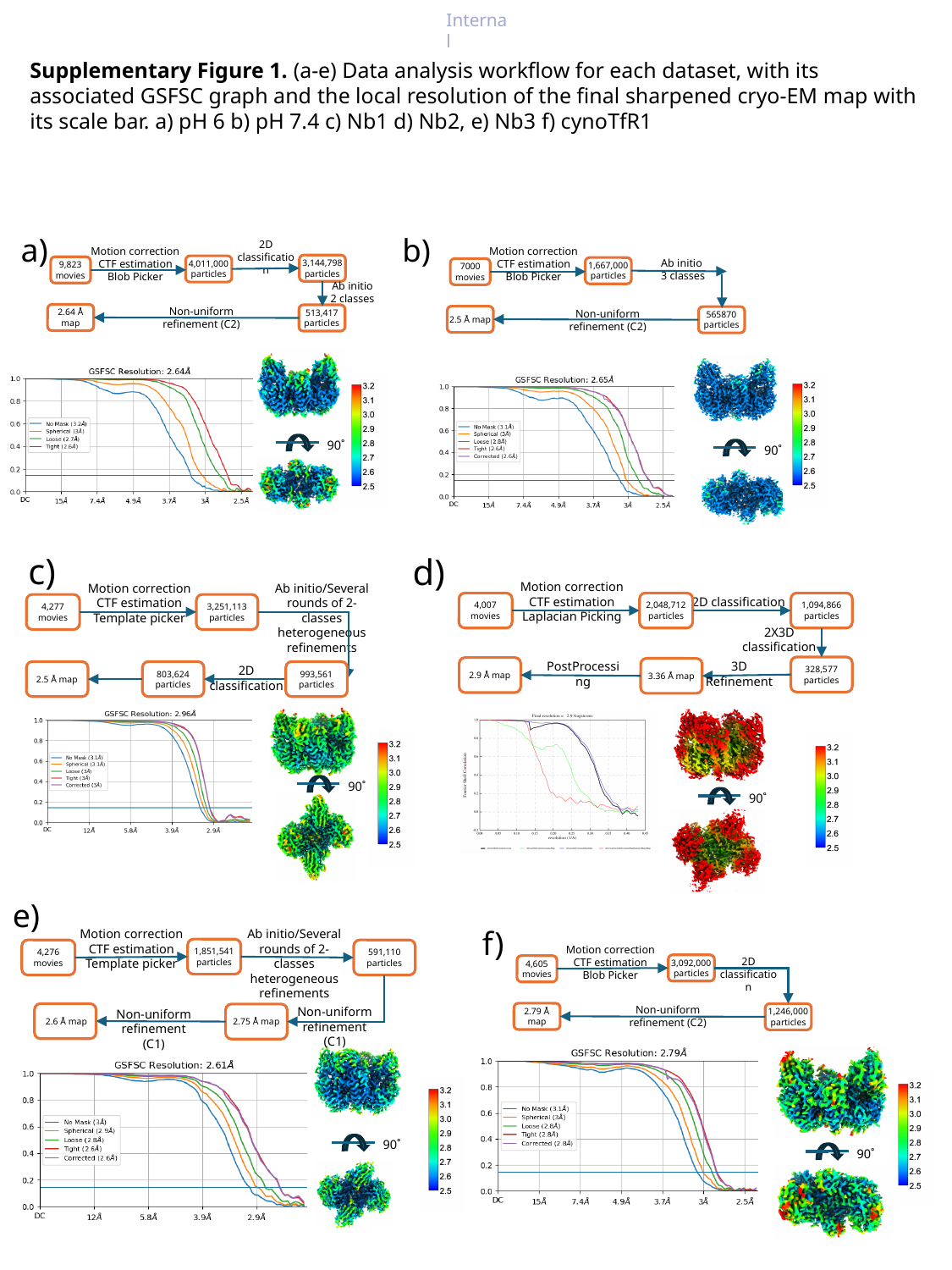

Supplementary Figure 1. (a-e) Data analysis workflow for each dataset, with its associated GSFSC graph and the local resolution of the final sharpened cryo-EM map with its scale bar. a) pH 6 b) pH 7.4 c) Nb1 d) Nb2, e) Nb3 f) cynoTfR1
# a)
b)
2D classification
Motion correction
CTF estimation
Blob Picker
Motion correction
CTF estimation
Blob Picker
Ab initio
3 classes
3,144,798
particles
4,011,000
particles
9,823
movies
1,667,000
particles
7000
movies
Ab initio
2 classes
Non-uniform refinement (C2)
Non-uniform refinement (C2)
2.64 Å map
513,417
particles
2.5 Å map
565870
particles
90˚
90˚
c)
d)
Motion correction
CTF estimation
Laplacian Picking
Motion correction
CTF estimation
Template picker
Ab initio/Several rounds of 2-classes heterogeneous refinements
2D classification
4,007
movies
2,048,712
particles
1,094,866
particles
4,277
movies
3,251,113
particles
2X3D classification
3D
Refinement
PostProcessing
2D classification
328,577
particles
2.9 Å map
3.36 Å map
2.5 Å map
803,624
particles
993,561
particles
90˚
90˚
e)
Motion correction
CTF estimation
Template picker
Ab initio/Several rounds of 2-classes heterogeneous refinements
f)
Motion correction
CTF estimation
Blob Picker
1,851,541 particles
4,276
movies
591,110
particles
2D classification
3,092,000
particles
4,605
movies
Non-uniform refinement (C2)
Non-uniform refinement (C1)
Non-uniform refinement (C1)
2.79 Å map
1,246,000
particles
 2.6 Å map
2.75 Å map
90˚
90˚

### Slide 4
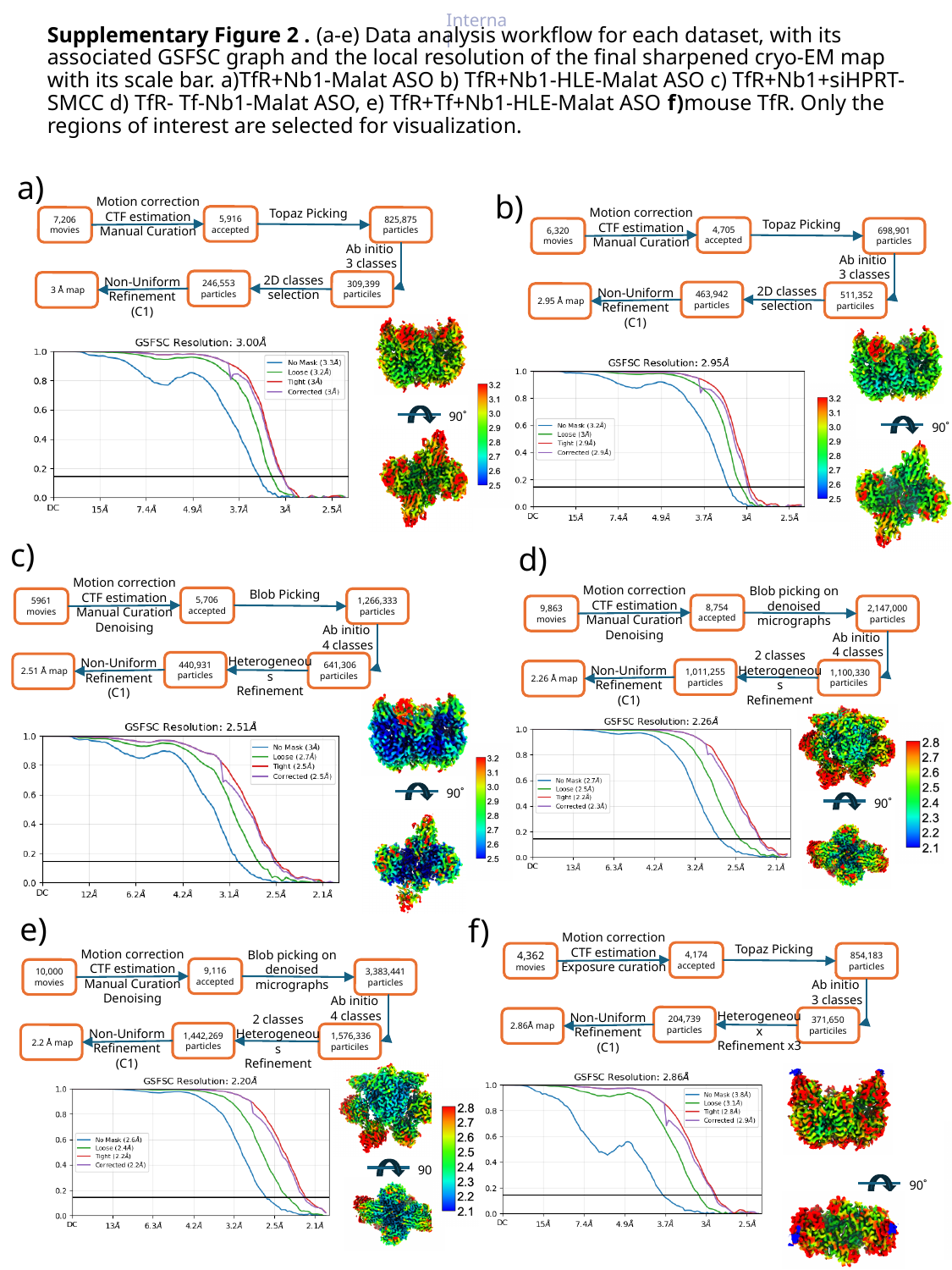

Supplementary Figure 2 . (a-e) Data analysis workflow for each dataset, with its associated GSFSC graph and the local resolution of the final sharpened cryo-EM map with its scale bar. a)TfR+Nb1-Malat ASO b) TfR+Nb1-HLE-Malat ASO c) TfR+Nb1+siHPRT-SMCC d) TfR- Tf-Nb1-Malat ASO, e) TfR+Tf+Nb1-HLE-Malat ASO f)mouse TfR. Only the regions of interest are selected for visualization.
# a)
b)
Motion correction
CTF estimation
Manual Curation
Motion correction
CTF estimation
Manual Curation
Topaz Picking
5,916
accepted
7,206
movies
825,875
particles
Topaz Picking
4,705
accepted
6,320
movies
698,901
particles
Ab initio
3 classes
Ab initio
3 classes
2D classes
selection
Non-Uniform
Refinement (C1)
246,553
particles
 309,399 particiles
 3 Å map
2D classes
selection
Non-Uniform
Refinement (C1)
463,942
particles
 511,352 particiles
 2.95 Å map
90˚
90˚
c)
d)
Motion correction
CTF estimation
Manual Curation
Denoising
Blob picking on denoised micrographs
8,754
accepted
9,863
movies
2,147,000
particles
Ab initio
4 classes
2 classes
Heterogeneous
Refinement
Non-Uniform
Refinement (C1)
1,011,255
particles
 1,100,330 particiles
 2.26 Å map
90˚
Motion correction
CTF estimation
Manual Curation
Denoising
Blob Picking
5,706
accepted
5961
movies
1,266,333
particles
Ab initio
4 classes
Heterogeneous
Refinement
Non-Uniform
Refinement (C1)
440,931
particles
 641,306 particiles
 2.51 Å map
90˚
e)
Motion correction
CTF estimation
Manual Curation
Denoising
Blob picking on denoised micrographs
9,116
accepted
10,000
movies
3,383,441
particles
Ab initio
4 classes
2 classes
Heterogeneous
Refinement
Non-Uniform
Refinement (C1)
1,442,269
particles
 1,576,336 particiles
 2.2 Å map
90˚
f)
Motion correction
CTF estimation
Exposure curation
Topaz Picking
4,174
accepted
4,362
movies
854,183
particles
Ab initio
3 classes
Heterogeneoux
Refinement x3
Non-Uniform
Refinement (C1)
204,739
particles
371,650
particiles
2.86Å map
90˚

### Slide 5
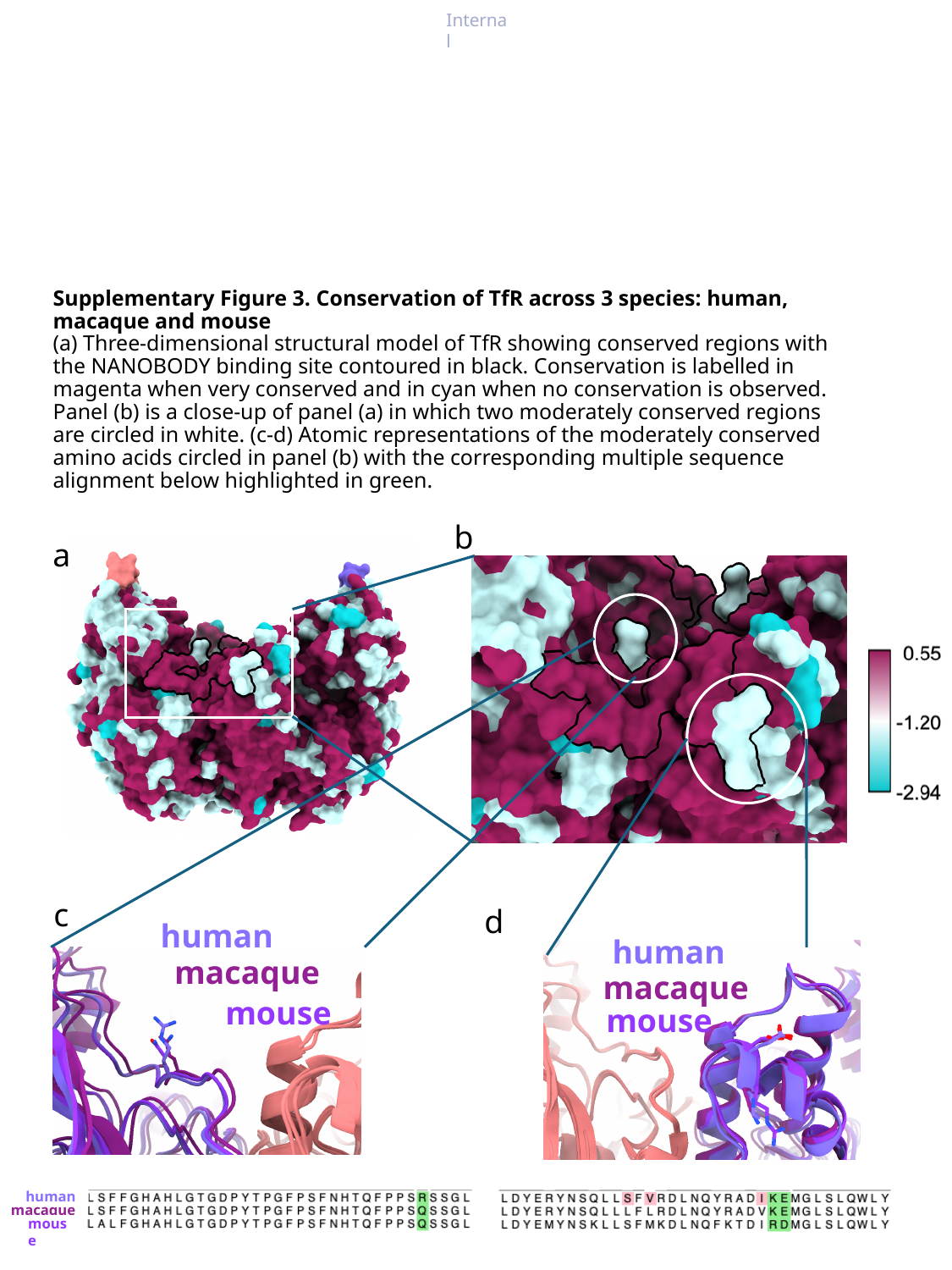

#### Supplementary Figure 3. Conservation of TfR across 3 species: human, macaque and mouse (a) Three-dimensional structural model of TfR showing conserved regions with the NANOBODY binding site contoured in black. Conservation is labelled in magenta when very conserved and in cyan when no conservation is observed. Panel (b) is a close-up of panel (a) in which two moderately conserved regions are circled in white. (c-d) Atomic representations of the moderately conserved amino acids circled in panel (b) with the corresponding multiple sequence alignment below highlighted in green.
b
a
c
d
human
human
macaque
macaque
mouse
mouse
human
macaque
mouse

### Slide 6
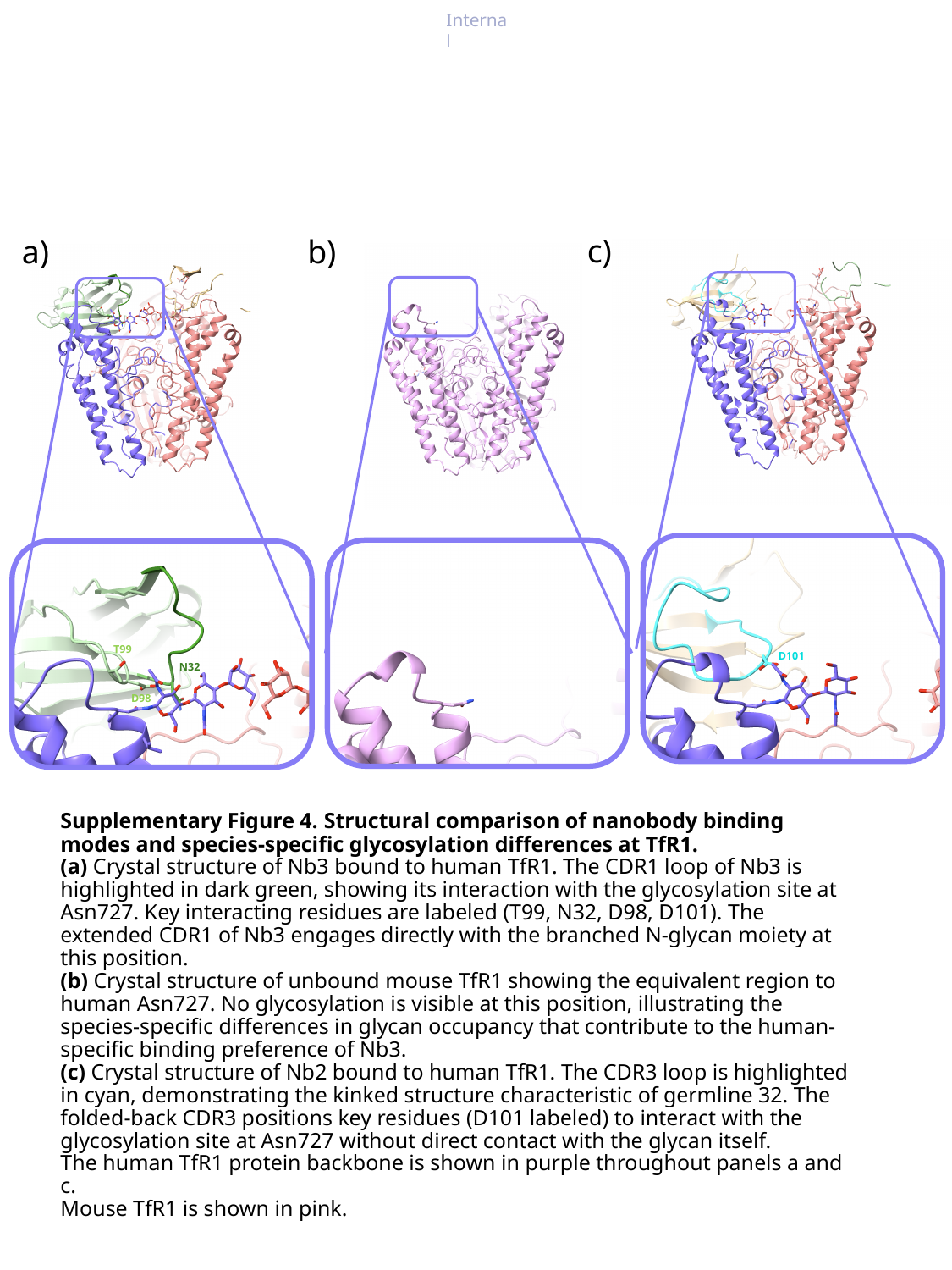

c)
D101
a)
T99
N32
D98
b)
#### Supplementary Figure 4. Structural comparison of nanobody binding modes and species-specific glycosylation differences at TfR1. (a) Crystal structure of Nb3 bound to human TfR1. The CDR1 loop of Nb3 is highlighted in dark green, showing its interaction with the glycosylation site at Asn727. Key interacting residues are labeled (T99, N32, D98, D101). The extended CDR1 of Nb3 engages directly with the branched N-glycan moiety at this position. (b) Crystal structure of unbound mouse TfR1 showing the equivalent region to human Asn727. No glycosylation is visible at this position, illustrating the species-specific differences in glycan occupancy that contribute to the human-specific binding preference of Nb3. (c) Crystal structure of Nb2 bound to human TfR1. The CDR3 loop is highlighted in cyan, demonstrating the kinked structure characteristic of germline 32. The folded-back CDR3 positions key residues (D101 labeled) to interact with the glycosylation site at Asn727 without direct contact with the glycan itself. The human TfR1 protein backbone is shown in purple throughout panels a and c. Mouse TfR1 is shown in pink.

### Slide 7
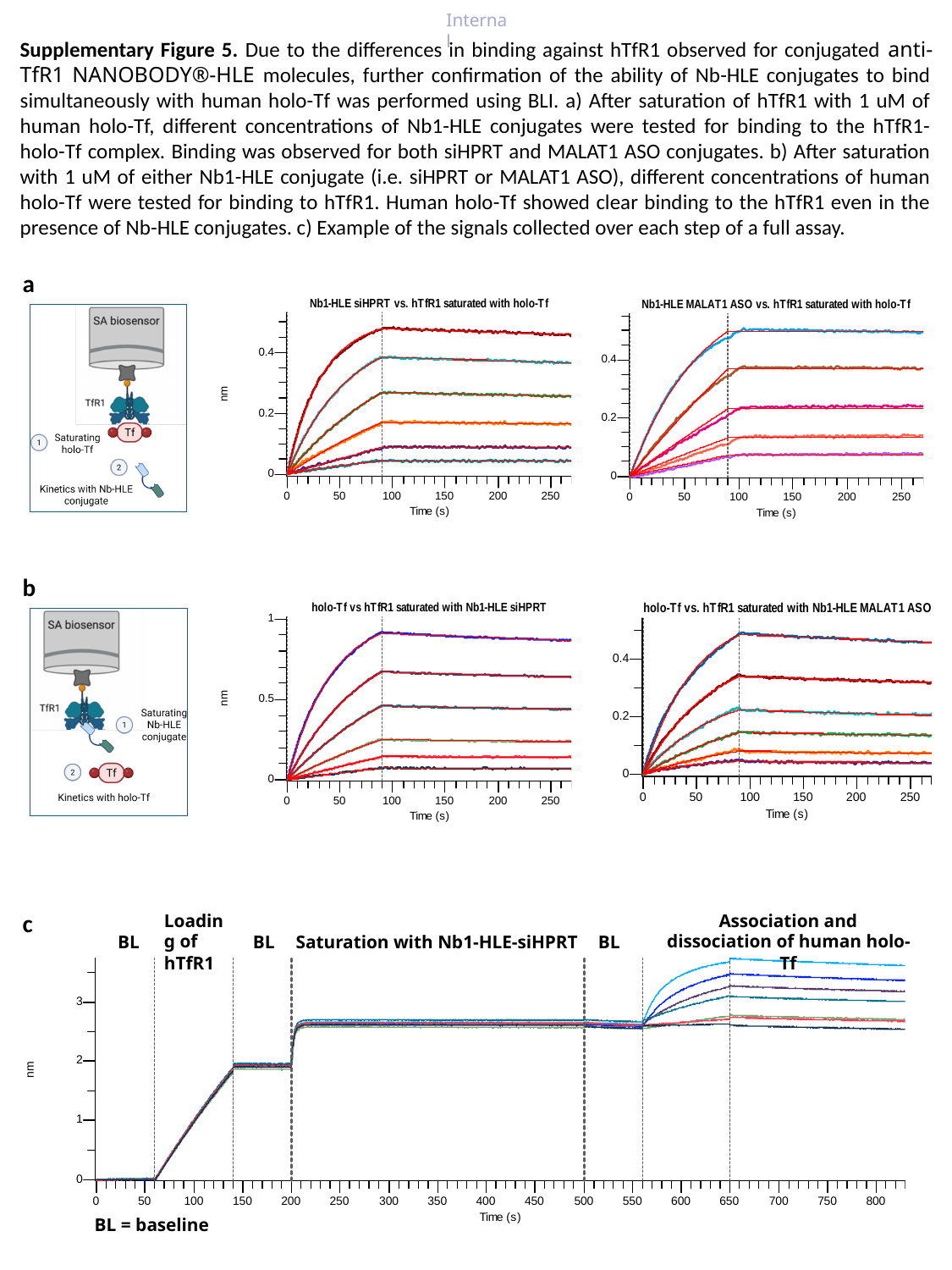

Supplementary Figure 5. Due to the differences in binding against hTfR1 observed for conjugated anti-TfR1 NANOBODY®-HLE molecules, further confirmation of the ability of Nb-HLE conjugates to bind simultaneously with human holo-Tf was performed using BLI. a) After saturation of hTfR1 with 1 uM of human holo-Tf, different concentrations of Nb1-HLE conjugates were tested for binding to the hTfR1-holo-Tf complex. Binding was observed for both siHPRT and MALAT1 ASO conjugates. b) After saturation with 1 uM of either Nb1-HLE conjugate (i.e. siHPRT or MALAT1 ASO), different concentrations of human holo-Tf were tested for binding to hTfR1. Human holo-Tf showed clear binding to the hTfR1 even in the presence of Nb-HLE conjugates. c) Example of the signals collected over each step of a full assay.
a
b
c
Loading of hTfR1
Association and dissociation of human holo-Tf
BL
BL
Saturation with Nb1-HLE-siHPRT
BL
BL = baseline

### Slide 8
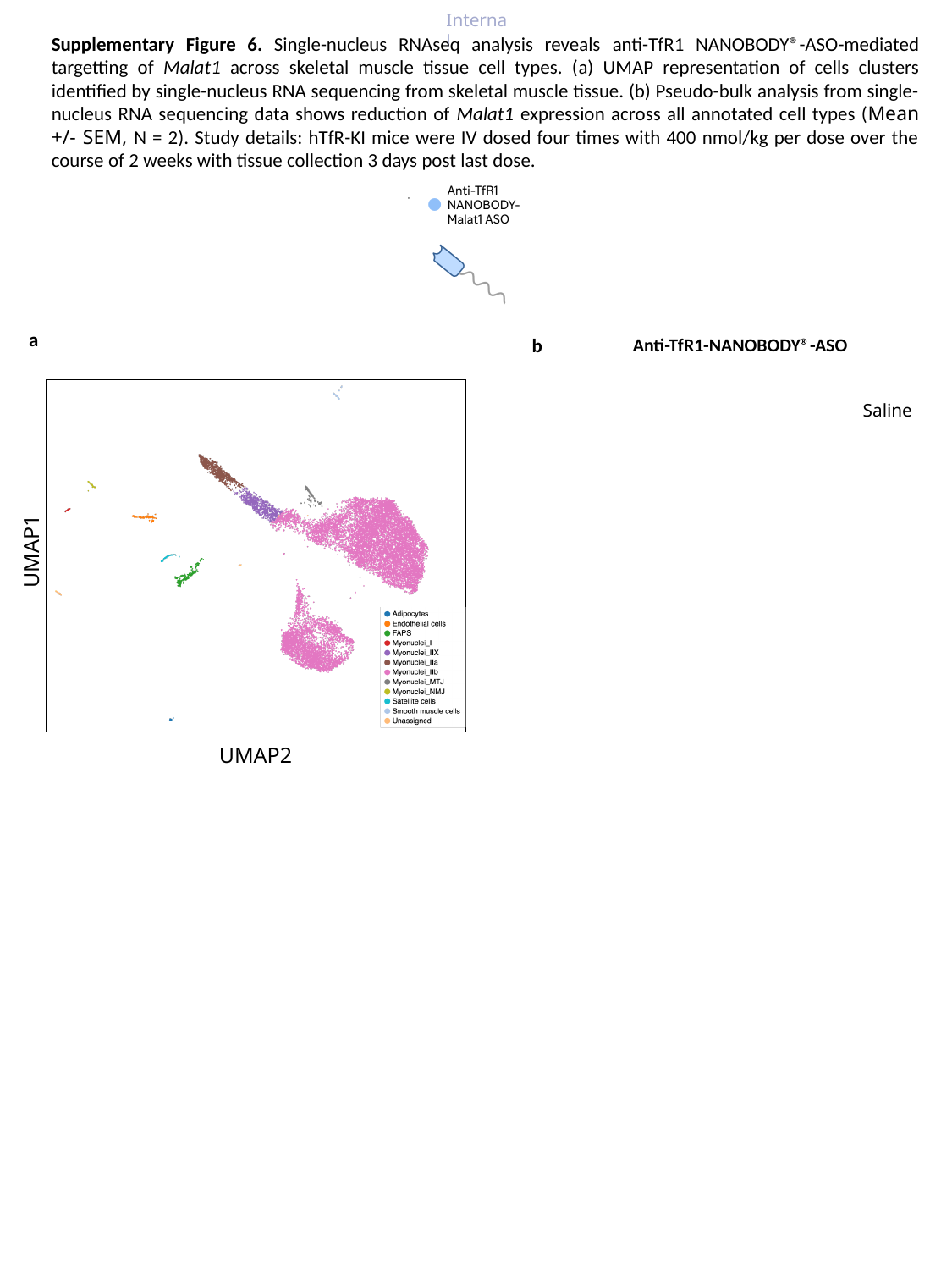

Supplementary Figure 6. Single-nucleus RNAseq analysis reveals anti-TfR1 NANOBODY®-ASO-mediated targetting of Malat1 across skeletal muscle tissue cell types. (a) UMAP representation of cells clusters identified by single-nucleus RNA sequencing from skeletal muscle tissue. (b) Pseudo-bulk analysis from single-nucleus RNA sequencing data shows reduction of Malat1 expression across all annotated cell types (Mean +/- SEM, N = 2). Study details: hTfR-KI mice were IV dosed four times with 400 nmol/kg per dose over the course of 2 weeks with tissue collection 3 days post last dose.
a
b
Anti-TfR1-NANOBODY®️-ASO
Saline
UMAP1
UMAP2

### Slide 9
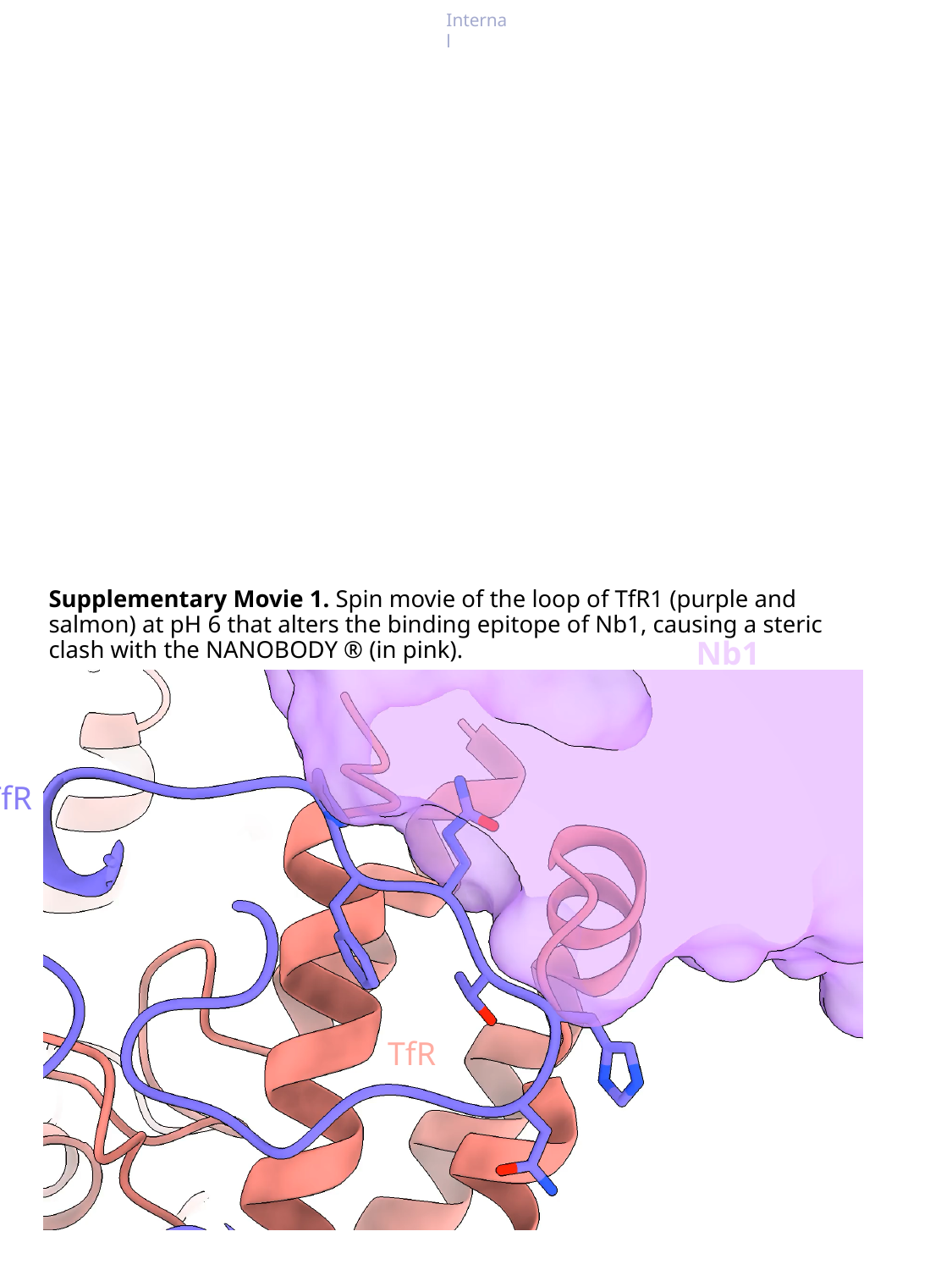

Supplementary Movie 1. Spin movie of the loop of TfR1 (purple and salmon) at pH 6 that alters the binding epitope of Nb1, causing a steric clash with the NANOBODY ® (in pink).
Nb1
TfR
TfR

### Slide 10
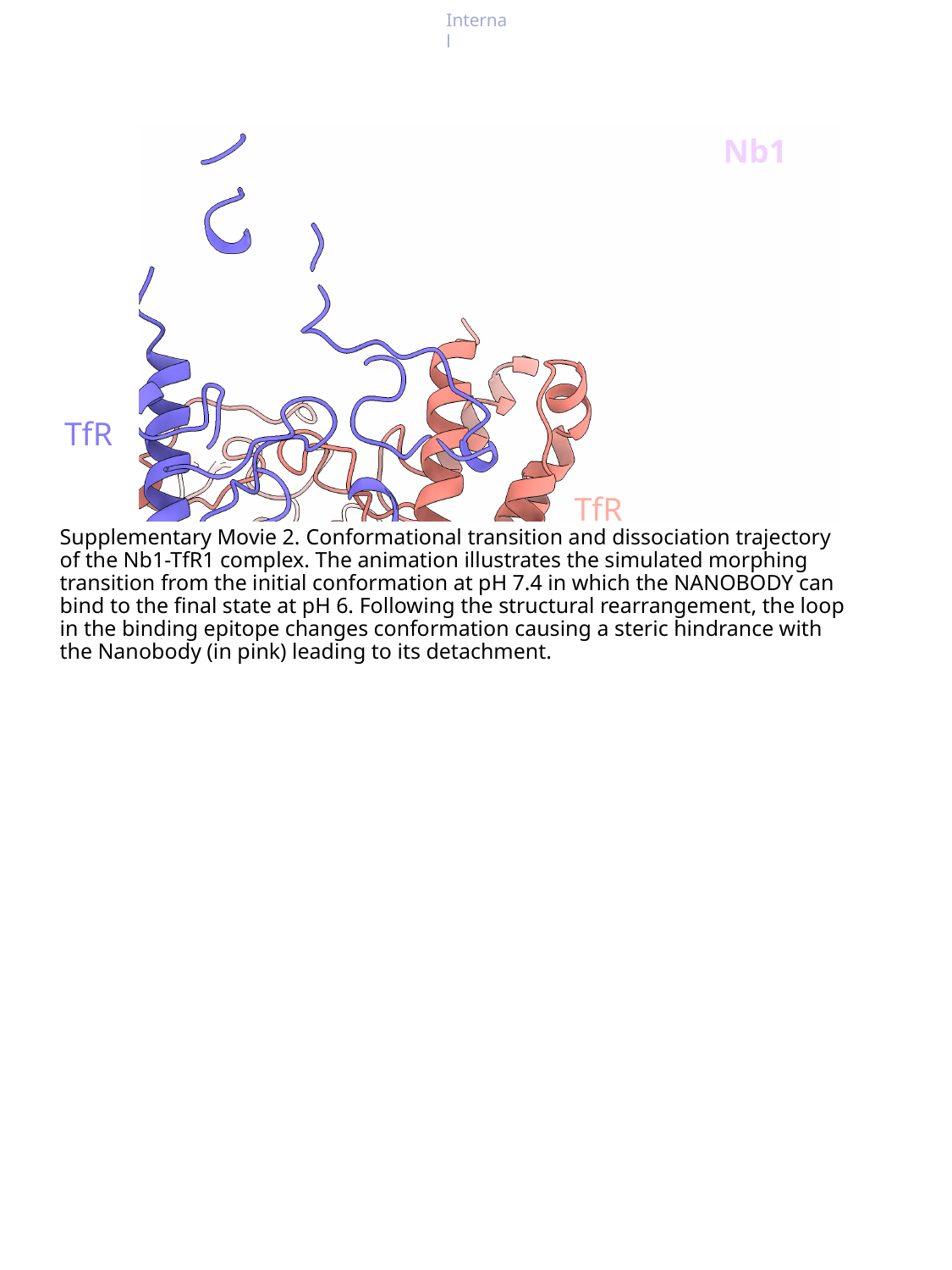

Nb1
TfR
TfR
Supplementary Movie 2. Conformational transition and dissociation trajectory of the Nb1-TfR1 complex. The animation illustrates the simulated morphing transition from the initial conformation at pH 7.4 in which the NANOBODY can bind to the final state at pH 6. Following the structural rearrangement, the loop in the binding epitope changes conformation causing a steric hindrance with the Nanobody (in pink) leading to its detachment.
